# Same visitors, different outcomes: floral phenotype gates pollinator-vectored nectar microbial establishment

**DOI:** 10.64898/2026.09.11.750984

**Authors:** Tueux Guillaume, Mumm Roland, Cankar Katarina, Jacquemot Marie-Pierre, Renard Océane, Langlade Nicolas Bernard, Carlier Aurélien

**Author notes:** Corresponding authors: Nicolas Langlade and Aurélien Carlier.

## Abstract

The microbiome of nectar has the potential to modify flower chemistry and plant-pollinator interactions. How its assembly differs between diurnal and nocturnal guilds, and how it depends on the flower they visit, remains poorly resolved in crops. We conducted field assays using two commercial sunflower cultivars (*Helianthus annuus*) that differ markedly in nectar volume and sugar composition, under four pollinator access treatments. Diurnal pollinator access, whether continuous or daytime-only, resulted in a cultivar-dependent microbiome signature. In one cultivar, nectar communities were characterized by the dominance of specialist taxa such as the yeast *Metschnikowia* and the bacterium *Acinetobacter*. However, neither taxa established dominance in the nectar of the other cultivar despite receiving substantial insect visits. Instead, continuous access alone lowered overall fungal diversity without restructuring composition, which was more strongly structured by inter-annual variation than by pollinator access treatment. Profiling of the volatile organic compounds in florets revealed clear differences between cultivars, but different access treatment induced only minor changes in the volatile blends. Our findings establish that pollinator guilds shape sunflower nectar microbiota in a genotype-dependent manner. We propose that diurnal insect visits seed nectar with specialist taxa, but whether those specialists establish depends on nectar availability and composition.

## INTRODUCTION

Floral nectar is not a sterile reward, but an exposed aerial habitat colonised by fungi and bacteria dispersed from the surrounding environment and by floral visitors [1]. Once established, these microbial communities are metabolically active: through sugar consumption and metabolites production, they shape the chemical composition of the nectar in which they grow [2]. Microbial metabolism may further affect the nutritional quality of the reward and, through it, pollinator health [3]. Such microbially driven changes may alter the floral volatile bouquet and, in turn, some of the cues that mediate plant– pollinator communication [4,5]. Whether such microbially driven shifts in volatile profiles arise under field conditions, and how they track visitor-mediated differences in microbiome composition, remains largely untested in crop systems.

The composition of nectar microbial communities is shaped by two processes: dispersal of propagules into the flower, and the ecological filters that determine community structure [1]. Floral visitors are major vectors of dispersal, carrying fungi and bacteria between flowers on their mouthparts and bodies, such that visitation can rapidly introduce new microbes [6]. Floral visitors are not the only inoculation route: wind, rain and direct contact with floral tissues also seed the nectar, contributing to microbial communities in those flowers that receive few or no visitors [6,7]. Microbes introduced to nectar then face a harsh environment: high osmotic pressure and plant secondary metabolites impose strong selection, so that only a few specialised taxa typically come to dominate [8,9]. Beyond this abiotic filtering, the order and timing of arrival may also shape community trajectories, because early arrivals may pre-empt resources and modify the chemical environment through priority effects [10].

Flower visitors often leave distinct microbial signatures in nectar [6,11]. For example, flowers accessible to both hummingbirds and bees harbour bacterial communities that are distinct from those accessible to bees alone [12]. Whether nocturnal visitors such as moths also leave microbial signatures and participate to microbe dispersal is far less studied [13]. The few studies addressing nocturnally visited flowers have compared visited and excluded plants on species that bloom exclusively at night [14], leaving unresolved the distinct contributions of day- and night-active guilds to a flower accessible to both. Yet diurnal and nocturnal visitors differ in foraging behaviour, body morphology and the resources they exploit, and may therefore deposit distinct microbial assemblages. Several studies have tested the contributions of different visitor guilds to pollen deposition through temporal partitioning of visitor access, for example by leaving flowers accessible only by day, only by night or not at all [15]. Applying the same partitioning to the nectar microbiome would allow each guild’s contribution to be assessed against a shared baseline of unvisited flowers, and against unrestricted access.

The outcome of dispersal and filtering is unlikely to be uniform across plants. Nectar volume itself has been associated with microbial establishment, with higher-volume flowers more likely to harbour bacteria [16]. In sunflower (*Helianthus annuus*), both nectar quantity and sucrose-to-hexose balance vary among genotypes, with consequences for pollinator attraction [17,18] as well as for the nectar fungal community [18]. Nectar traits may therefore modulate how visitation influences nectar communities, both through pollinator attractiveness and through ecological filters affected by nectar chemistry.

Because nectar chemistry and secretion are variable between genotypes, sunflower offers a suitable system to test how floral traits modulate the contribution of diurnal and nocturnal visitors to nectar microbiome assembly. We conducted a field pollinator-exclusion experiment over two consecutive growing seasons on two commercial cultivars that contrast in nectar volume and sugar composition. Individual plants were assigned to one of four 24-h treatments: no visitor access, continuous access, daytime-only access or nighttime-only access. This design allowed us to test three hypotheses: (i) nectar community structure and/or composition depends on floral context, producing contrasting community trajectories between cultivars under the same access treatment; (ii) diurnal and nocturnal guilds contribute unequally to nectar microbiome assembly; (iii) visitor-mediated microbial differences extend profiles of floral volatile compounds.

## MATERIALS AND METHODS

### Sunflower culture and phenotyping

Plants were grown at the INRAE experimental station in Auzeville-Tolosane, France, during the 2024 and 2025 field seasons. Two high-oleic sunflower cultivars were used: Celesto (Syngenta) and Idillic (LIDEA). Trials were conducted in July for both years. Pollinator access was controlled using mesh pollination bags (Vilutis and Co.). At the start of the experiment at 06:00 (UTC+2), plants were assigned to one of four pollinator-access treatments lasting 24 h, until the next day at 06:00: continuously netted (no access), fully unnetted (continuous access), unnetted until 22:00, then netted (daytime access), or netted until 22:00 then unnetted (nighttime access), a transition time corresponding to local sunset at this latitude in July (Figure 1A).

**Figure 1.**
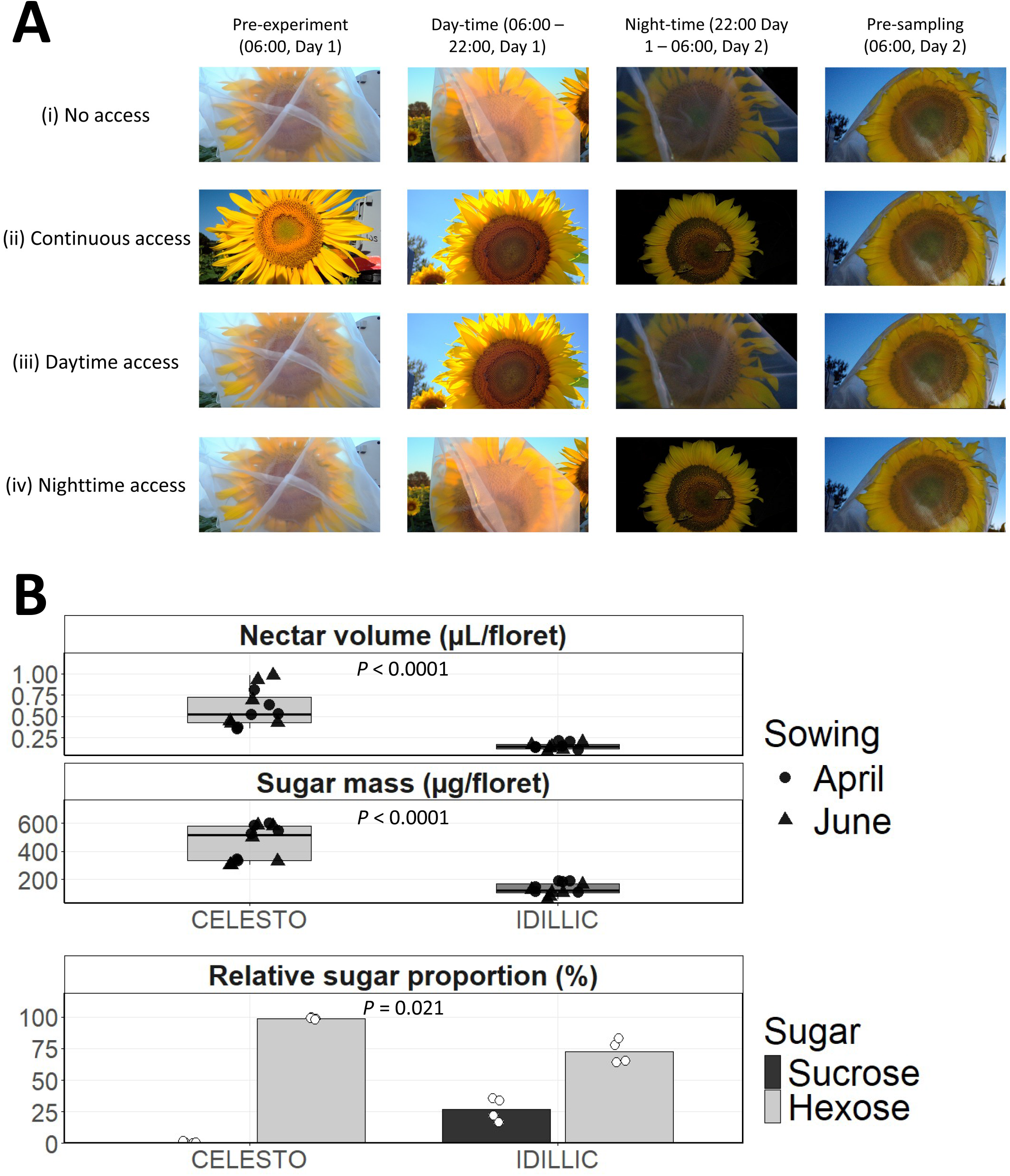
Floral traits and experimental design. (A) Experimental design showing the four pollinator exclusion treatments. Each row represents a treatment: (i) No pollinators access, (ii) continuous pollinators access, (iii) Daytime pollinators access only (netted at night), and (iv) Nighttime pollinators access only (netted during the day). Columns show the state of each plant at four time points across the 24-h experiment. Plants assigned to the no access, daytime access and nighttime access treatments were enclosed in mesh bags before treatment assignment at 06:00 (Day 1). Continuous-access plants remained non-netted throughout. All sampled florets were at the same developmental stage and had received the same treatment duration. Photographs were taken at the INRAE experimental station in Auzeville-Tolosane, France. (B) Nectar volume (µL per floret), sugar mass (µg per floret), and relative sugar composition of Idillic and Celesto sunflower cultivars. For nectar volume and sugar mass, each point represents the mean value of one plant (n = 12 per cultivar), with shape indicating sowing date. *P*-values indicate results of Wilcoxon rank-sum tests comparing cultivars. For sugar composition, points represent individual plant means (n = 4 per cultivar) for sucrose and hexose (glucose + fructose) proportions. Hexose proportions were compared between cultivars with a Kruskal-Wallis test.

### Floral traits

Nectar was harvested from six florets per head using 2-µL glass microcapillaries (Hirschmann, Germany), approximately 24 h after anthesis at the pistillate stage (10:00– 13:00 local time). Sampling was conducted in 2025 across two sowings in the same field, established in April and June and sampled at flowering in July and August, respectively, with 6 plants per cultivar sampled per sowing. Nectar volume was recorded from each capillary, and soluble-sugar concentration (°Bx) was measured with a refractometer (Bellingham & Stanley, UK). Sugar mass per floret was calculated with the formula detailed by Cruden and Hermann [19]. Sucrose, glucose and fructose concentrations were determined enzymatically (Megazyme K-SUFRG kit) for four plants per cultivar from a single sowing, with plant-level means pooled across florets. Nectar volume and sugar mass, with plant-level means averaged across florets, were compared between cultivars and between sowings with Wilcoxon rank-sum tests, and hexose proportion (glucose + fructose) between cultivars with a Kruskal-Wallis test. Capillary-level measurements are available in the data repository (https://doi.org/10.57745/LGKSOT).

### Quantification of pollinator visits

Insect visitation was quantified from time-lapse images acquired every 30 seconds throughout the experiment using Wingscapes TimelapseCam Pro cameras. Insects were detected and classified into three categories: non-bombus bees (*Apis mellifera*, *Megachile spp*., Halictidae) hereafter referred to as “bees”, bumblebees and moths using an internally developed model [20] based on the YOLO11x architecture (https://forge.inrae.fr/astr/public/pollicrop, model 09-25). Model performance was validated against expert annotations on a randomly selected subset of 5,000 images, with per-class precision, recall and F1 scores reported in Table S1 and raw detection counts per image in the data repository (https://doi.org/10.57745/LGKSOT).

Camera monitoring and nectar microbiota sequencing targeted the same plants, but the two datasets do not fully overlap because of technical issues with nectar sampling or the cameras. Sample sizes for each are reported separately in Table S2.

Consistency between continuous access and single-window treatments was assessed separately by cultivar, comparing diurnal (bee, bumblebee) and nocturnal (moth) visit rates within the corresponding time window of continuous-access plants to the matching daytime- or nighttime-access treatment, using Wilcoxon rank-sum tests with Benjamini–Hochberg correction across the six tests.

### PCR and sequencing of nectar microbiota samples

Flowers were re-bagged at the end of the 24-h access window (06:00) and remained bagged until nectar was collected. Nectar samples were collected in 2024 and 2025, 24 h post-anthesis at the pistillate stage (09:00–12:00 local time) by pooling nectar from several florets from the same head into six sterile 2 µL (Idillic) or 5 µL (Celesto) glass microcapillaries (Hirschmann). Only inflorescences with detectable nectar were retained for microbiota sampling, yielding 109 plants sampled across cultivars, pollinator access treatments and trial years (Table S2). Nitrile gloves were sanitised with 70% ethanol between plants. Sampling controls included empty capillaries briefly exposed to air, and touched with gloved fingertips. Extraction blanks, in which water was processed through the full DNA extraction. Capillaries were stored at −80 °C in sterile 2 mL tubes before DNA extraction (DNeasy PowerSoil Pro kit, QIAGEN). Negative controls were processed alongside nectar samples, and a positive control (ZymoBIOMICS Microbial Community Standard, Zymo Research) was carried through all downstream steps.

The near full-length bacterial 16S rRNA gene and the fungal ITS1–ITS4 region were amplified with adapter-tailed primers 16SF/16SR and ITS1F/ITS4R, respectively (Table S3) as described previously [18]. Sequencing was performed on a PromethION 2 with R10.4.1 flow cells, and POD5 signals were basecalled with Dorado v0.9 (dna_r10.4.1_e8.2_400bps_sup@v5.0.0 model; --no-trim and --barcode-both-ends options enabled).

Raw sequencing reads were processed as described previously [18], using custom Python scripts available at https://forge.inrae.fr/aurelien.carlier/ont-metabarcoding. Briefly, sequences were clustered into OTUs at 97% identity with VSEARCH [21], and chimeras and sequencing artefacts were removed with VSEARCH UChime [22] and mumu [23] with default settings. Taxonomy was assigned in QIIME 2 v.2023.9.15 [24] with the VSEARCH classifier set at an 80% identity threshold, using the SILVA 138 NR99 database for 16S sequences and the UNITE NR 99 database for fungal ITS. Raw sequencing data for all samples are deposited at the European Nucleotide Archive under study accession PRJEB121466.

### 16S Microbial diversity analysis

Validation of the sequencing and analysis procedures was assessed via recovery of OTUs corresponding to species present in the mock community sample. OTUs exceeding 5% relative abundance in at least one extraction blank were removed prior to analysis, except chloroplast-assigned OTUs, as described previously [18]. OTUs with fewer than 10 total reads and samples with fewer than 1,000 reads were discarded. The taxonomic assignment of *Acinetobacter* OTUs was performed by submitting the near full-length representative sequence (1459 bp) of each cluster to the EzBioCloud 16S-based ID online service [25].

### ITS Microbial diversity analysis

Non-fungal (plant) OTUs were discarded based on Kingdom assignment. OTUs reaching more than 5% of the reads in any extraction blank were removed from the entire dataset, while OTUs exceeding 5% in any sampling blank were set to zero only in samples from the same field season. OTUs assigned to the class Malasseziomycetes, common skin-associated commensals likely introduced during handling, were also excluded. Initial OTU clustering at 97% similarity followed by curation with *mumu* resulted in substantial oversplitting within the genus *Metschnikowia*, yielding 184 OTUs assigned to this genus. There is high intra- and inter-genomic diversity of ITS variability in this clade [26], which prevents straightforward threshold-based resolution. To address this and avoid inflating diversity estimates, a phylogeny-informed merging procedure was applied for all OTUs assigned to the genus *Metschnikowia*. 184 OTU sequences together with 169 ITS reference sequences of *Metschnikowia* and related genera (*Clavispora*, *Kodamaea*) downloaded from the UNITE database were selected (data repository; https://doi.org/10.57745/LGKSOT), and aligned with MAFFT v7.505 [27]. The alignment was trimmed with Trimal [28] to remove columns with >95% of gap characters. A Maximum Likelihood phylogeny was inferred in FastTree2 using the -gtr and -gamma parameters [29] and rooted at the midpoint The clade containing all *Metschnikowia reukaufii* reference sequences, the dominant yeast species consistently reported in floral nectar [30], was expanded node-by-node, moving toward the root, until it first included a reference sequence from another named species. The merging threshold was set to the maximum pairwise patristic distance among OTUs in the clade just before that point, plus a negligible epsilon to avoid ties at the boundary. Clades containing two or more OTUs with a maximum pairwise distance below this threshold were then merged, starting with the largest clades, excluding any OTU already assigned to a larger cluster, to produce a non-overlapping set of merge groups (Figure S1). This reduced the 184 putative *Metschnikowia* OTUs to 8 high confidence OTUs. Read counts of merged OTUs were summed to produce the final feature table. OTUs with fewer than 10 total reads and samples with fewer than 1,000 reads were excluded, yielding a final dataset of 104 samples and 357 OTUs.

Alpha diversity was estimated as coverage-standardised Hill numbers at 95% sample coverage using iNEXT [31]. We report richness (Hill q = 0), the exponential of Shannon entropy (Hill q = 1; ENS1) and inverse Simpson (Hill q = 2; ENS2). Richness was square-root transformed, and ENS1, ENS2 were log-transformed prior to analysis to meet model assumptions. Each metric was analysed by ANOVA with treatment, cultivar, their interaction, and trial year as fixed effects, followed by Tukey-adjusted pairwise contrasts between treatments within each cultivar from estimated marginal means. Fungal alpha diversity analyses were additionally run on no-access samples alone, with cultivar and trial year as factors.

### Beta diversity analyses

Beta diversity was assessed using four complementary dissimilarity metrics applied to both 16S and ITS datasets: Bray–Curtis on relative abundances, Jaccard on presence–absence data, and unweighted and weighted UniFrac incorporating phylogenetic distances.

Analyses were conducted separately for each cultivar. Differences in community composition between visitor treatments were tested by PERMANOVA (adonis2, 99,999 permutations; vegan) with treatment and trial year included as model terms and tested using marginal (type III) sums of squares. Homogeneity of multivariate dispersions was verified beforehand using betadisper and permutest (9,999 permutations). PERMANOVA outputs were interpreted only where dispersion was homogeneous. Ordinations were obtained by principal coordinates analysis (PCoA) with 95% confidence ellipses. Outputs for all four metrics, for both 16S and ITS, are provided in Table S4. Pairwise comparisons between treatments were performed with Benjamini–Hochberg correction applied across the six pairwise comparisons within each metric, using 9,999 permutations.

### Isolation of microbes from nectar samples

Nectar samples were collected 24 h post-anthesis at the pistillate stage, pooling nectar from several florets per head into sterile glass microcapillaries), from a separate set of plants. Flowers were re-bagged at the end of the 24-h access window (06:00) and remained bagged until nectar was collected. Nectar samples were resuspended in 500 µL of sterile phosphate-buffered saline (PBS) and spread onto tryptic soy agar (TSA), R2A agar, potato dextrose agar (PDA) and yeast-malt (YM) agar, then incubated at 28 °C. As nectar volume was not measured prior to resuspension, growth scores reflect microbial load per sample rather than per unit volume of nectar. Because reliable CFU counts were not feasible at high colony densities, each plate was assigned a semi-quantitative growth score: 0 (no colonies), 1 (1-10), 2 (11-50), 3 (51-500) and 4 (>500 colonies; Figure S2). Plates were scored by the same observer, 6 days after sampling in 2024 and at 4 days in 2025. For each sample, growth scores were averaged across the culture media, and pairwise comparisons between pollinator access treatments were performed on these per-sample means with Wilcoxon rank-sum tests and Benjamini-Hochberg correction (n = 6-7 per treatment in Celesto; n = 3-7 per treatment in Idillic; Table S5). Raw culture density data (both seasons) and pictures of plates (2025 season only) are provided in the data repository (https://doi.org/10.57745/LGKSOT).

### Pollinator visitation and fungal microbiome relationships

Visitation data from time-lapse images were summarised per plant as mean number of visits per photograph for three insect categories (bees, bumblebees, moths). Analyses were focused on treatments and time periods at which flowers accessible to insects (continuous, daytime and nighttime access; n = 67 plants with both usable camera data and a retained ITS profile). Bagged plants were not monitored. To test whether visit intensity predicted microbiome structure beyond the categorical effect of visitation treatment, diurnal (bee + bumblebee) and nocturnal (moth) visit rates were analysed within paired treatment windows (daytime access + continuous access for diurnal rate, nighttime access + continuous access for nocturnal rate), separately by cultivar. For continuous-access plants, diurnal and nocturnal visit rates were each computed using only the subset of photographs falling within the corresponding window, to avoid dilution by the complementary period’s near-zero counts. Alpha diversity measures (Hill numbers q = 0, 1, 2) were modelled by linear regression including visitation treatment, visit rate (diurnal or nocturnal, as appropriate) and trial year as covariates. Beta-diversity was tested analogously, regressing each of the four-dissimilarity metrics on visit rate with trial year as covariate (PERMANOVA, marginal terms, 99,999 permutations).

### Analysis of volatile organic compounds (VOCs) of florets

Volatile profiling was performed on seven plants per cultivar × access-treatment combination (56 plants in total). These plants were sampled from the same set of those used for nectar microbial isolation, but were distinct from those used for microbiota sequencing. Florets were collected at the pistillate stage, 24 h after pollen presentation. For each sample, ten to fifteen florets were immediately snap-frozen in liquid nitrogen and stored at -80 °C until processing. Tissue was homogenized using a Retsch MM400 mixer mill with a pre-cooled 10 mL stainless steel grinding jar (Qiagen), re-cooled between cycles with liquid nitrogen to maintain sample integrity. VOCs were analysed by solid-phase microextraction coupled with gas chromatography mass spectrometry (SPME GC-MS) as described previously [32]. Briefly, 0.02g frozen floret powder was mixed 1mL of a 5M CaCl_2_-EDTA solution (pH 7.5; Sigma-Aldrich, The Netherlands) in a pre-cooled 10mL glass headspace vial (BGB, Germany). Vials were closed with magnetic screw caps (8mm hole) with Silicone/PTFE septa (BGB, Germany). A pooled mixture of powders from 27 samples was prepared, and 0.02 g aliquots were used as quality-control (QC) samples and analysed alongside the biological samples. A homologous series of n-alkanes (C_6_ – C_21_; Sigma-Aldrich, the Netherlands), was analysed using the same SPME GC-MS method to calculate retention indices (RIs).

Samples were incubated at 50 °C for 15 min with agitation at 250 rpm. Headspace volatiles were then extracted for 15 min at 50 °C using a 50/30 µm PDMS/DVB/CAR SPME fibre (Supelco, PA, USA) with an MPS-2 autosampler (Gerstel, Germany). Analytes were transferred to an Agilent GC7890A gas chromatograph coupled to a 5975C quadrupole mass spectrometer by thermal desorption in a Gerstel CIS4 at 250 °C for 2 minutes in splitless mode, under a constant helium flow of 1ml/min. The column used was a Zebron ZB-5MSplus (30 m x 0.25 mm i.d. x 1.00 μm film thickness; Phenomenex, the Netherlands). The GC oven was maintained at 45 °C for 2 min, heated to 230 °C at 5 °C/min, and then heated to 280 °C at 25 °C/min and held for 2 min. The column effluent was ionised by electron impact at 70 eV and ions were scanned over an m/z range of 33–330. A solvent delay of 3 minutes was applied.

Raw data were processed following an metabolomics workflow centered on the MetAlign and MSClust software packages as previously described [32]. Following baseline correction, peak picking and alignment of the mass signals, mass features detected more than three biological replicates within each sample group were retained. VOCs were identified by matching the reconstructed mass spectra and calculated RIs with those of authentic reference standards and with spectra using the NIST20 Mass Spectral library and in-house databases.

Peak areas were log10(x + 1)-transformed. On the 56 plants sampled, one sample was lost during tissue homogenisation. One further sample was excluded as an outlier following visual inspection of the ordination. For a total of 111 GC-MS detected VOCs, Bray–Curtis dissimilarities were calculated on relative abundances. Cultivar effect was tested by PERMANOVA (adonis2, 9,999 permutations). Treatment effects were then tested separately per cultivar. Dispersion homogeneity was verified with betadisper and permutest (9,999 permutations). Ordinations used PCoA with 95% confidence ellipses. For each of the 111 compounds, linear models were used to test the effect of access treatment across all four treatment levels (ANOVA). *P*-values for the treatment effect were Benjamini–Hochberg-adjusted within each cultivar; q < 0.05 was considered significant. For compounds with a significant treatment effect, Šidák-adjusted pairwise comparisons between modalities were extracted via emmeans and summarised as compact letter groupings.

### Statistical software and packages

All analyses were performed in R v4.5.1 using RStudio 2023.09.1. Data import and manipulation relied on the tidyverse packages dplyr v1.1.4, tidyr v1.3.1, purrr v1.0.4, tibble v3.3.0, readr v2.1.6, readxl v1.4.5, forcats v1.0.1 and lubridate v1.9.4. Microbial community and diversity analyses used phyloseq v1.52.0, vegan v2.7.1, iNEXT v3.0.1 and pairwiseAdonis v0.4.1. Phylogenetic analyses and visualization used ape v5.8.1, phytools v2.5.2, ggtree v3.16.3 and ggtreeExtra v1.18.1. Linear models and post-hoc comparisons were performed with car v3.1.3, emmeans v2.0.0, multcomp v1.4.29, multcompView v0.1.10 and broom v1.0.10. Figures were produced with ggplot2 v4.0.3, ggtext v0.2.0, ggnewscale v0.5.2, ggh4x v0.3.1, patchwork v1.3.2, pheatmap v1.0.13 and scales v1.4.0.

## RESULTS

To test how diurnal and nocturnal pollinator guilds shape nectar microbiome assembly, and whether floral context modulates this effect, we conducted a two-year field exclusion experiment on two sunflower cultivars contrasting in nectar volume and sugar composition (Celesto, Idillic). Individual plants were assigned to one of four 24-h pollinator access treatments: no access, continuous access, daytime access or nighttime access and then monitored for visitor activity, nectar microbiome composition, microbial load and floral volatiles.

### Contrasting nectar volumes and composition between the Celesto and Idillic cultivars

Celesto and Idillic are two high-oleic sunflower cultivars commonly grown for oilseed production. Flowers of the Celesto cultivar produced substantially more nectar than Idillic, both in volume (0.59 ± 0.21 vs 0.15 ± 0.04 µL per floret, Wilcoxon rank-sum, *P* < 0.0001) and sugar mass per floret (463 ± 126 vs 133 ± 44 µg, *P* < 0.0001), representing approximately a 3.5- to 4-fold difference in both traits (Figure 1B). Sowing date (April vs June 2025 sowings) had no effect on either variable (volume: *P* = 0.95; sugar mass: *P* = 0.26). Nectar sugar composition also differed markedly between cultivars: Celesto nectar was hexose-rich (glucose + fructose: 99.0 ± 0.8%), while Idillic nectar was composed of 72.8 ± 9.3% hexoses and 27.2 ± 9.3% sucrose (Figure 1B, Table S6).

### Netting flowers during day or night effectively partitioned visitor guilds

The contrasting nectar phenotypes of the Celesto and Idillic sunflower cultivars offer a valuable experimental system to understand how insect visits shape nectar microbiome assembly. Monitoring of insect visits using trap cameras on the plants that were sampled for nectar microbiome, showed that daytime-access plants received predominantly bee visits (1.34 detections/photo on average for the Celesto cultivar vs. 0.379 for Idillic), with moderate bumblebee activity (0.069 in Celesto; 0.049 in Idillic), and negligible moth activity (≤0.006 in both cultivars) (Figure 2). Nighttime-access plants were visited almost exclusively by moths (0.203 detections/photo in Celesto; 0.206 in Idillic), with low mean rates of bees (≤0.011 detections/photo) and bumblebees (≤0.007 detections/photo) (Figure 2). Continuous-access plants received all three guilds. Diurnal visit rates were lower under continuous access than daytime access in Celesto (bee: Wilcoxon rank-sum test, BH-adjusted, q = 0.036; bumblebee: q = 0.006), but not in Idillic (q = 0.341, 0.256). Nocturnal rates did not differ between continuous and nighttime access in either cultivar (q = 0.946, 0.253).

**Figure 2.**
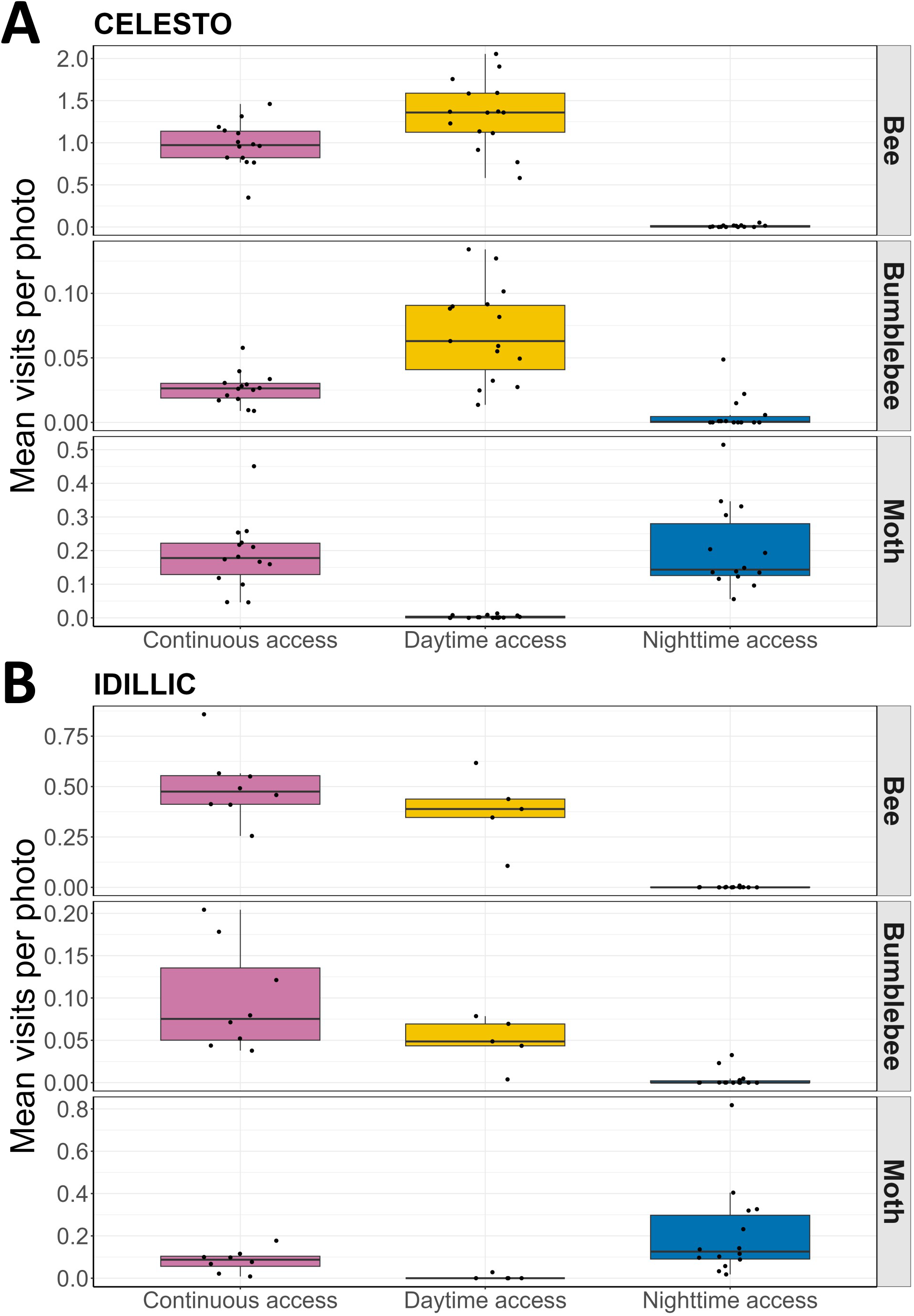
Pollinator visit rates per plant across treatments and cultivars. Mean number of detections per photo per plant for bees, bumblebees, and moths, recorded in the continuous access, daytime access, and nighttime access treatments, for (A) Celesto and (B) Idillic. For continuous access, bee and bumblebee rates were computed from photographs taken before 22:00 and moth rates from photographs taken from 22:00 onward, matching the time windows used for the daytime- and nighttime-access treatments. Each point represents one plant. Detections were classified using a YOLO11x-based model (PolliCrop). The no access treatment was excluded, as no camera monitoring was performed

Together, these treatments effectively decoupled diurnal and nocturnal visitor guilds within two cultivars that differ markedly in nectar volume and sugar composition, providing the experimental basis to test whether guild identity and floral context shape nectar microbiome assembly.

### Contrasting nectar bacterial community composition in sunflower cultivars

We first asked whether this visitor partitioning was reflected in nectar bacterial community composition. After filtering contaminant OTUs identified in extraction blanks, the 108 16S amplicon samples were represented by an average of 49,641 reads per sample and 9.2 OTUs per sample. A single OTU assigned to the genus *Acinetobacter* dominated nectar communities of the Celesto cultivar, and accounted on average for 98.6% and 99.8% of 16S reads in continuous-access and daytime-access samples, respectively (Figure S3). In contrast, chloroplast-derived sequences from *Helianthus annuus* (Table S7) dominated in the no-access and nighttime-access treatments, accounting on average for 100% and 92.1% of 16S reads, respectively and indicated the very low abundance of bacteria in these samples. Bacterial reads represented a small fraction of the 16S dataset in Idillic (0.1–7.3%). However, the same OTU assigned to *Acinetobacter* was also present in 7 out of 13 continuous- and daytime-access Idillic nectar samples, although at lower relative abundance ranging from 0.2% to 28.8% (continuous-access, n = 4) and from 0.02% to 2.8% (daytime-access, n = 3) among the samples with detection (Figure 3A). The representative near full-length 16S rRNA sequence of the OTU displayed 99.5% identity to that *Acinetobacter pollinis* SCC77^T^, and 99.31% identity to *Acinetobacter nectaris* CIP 110549^T^, two strains isolated from honey bees and floral nectar, respectively [33].

**Figure 3.**
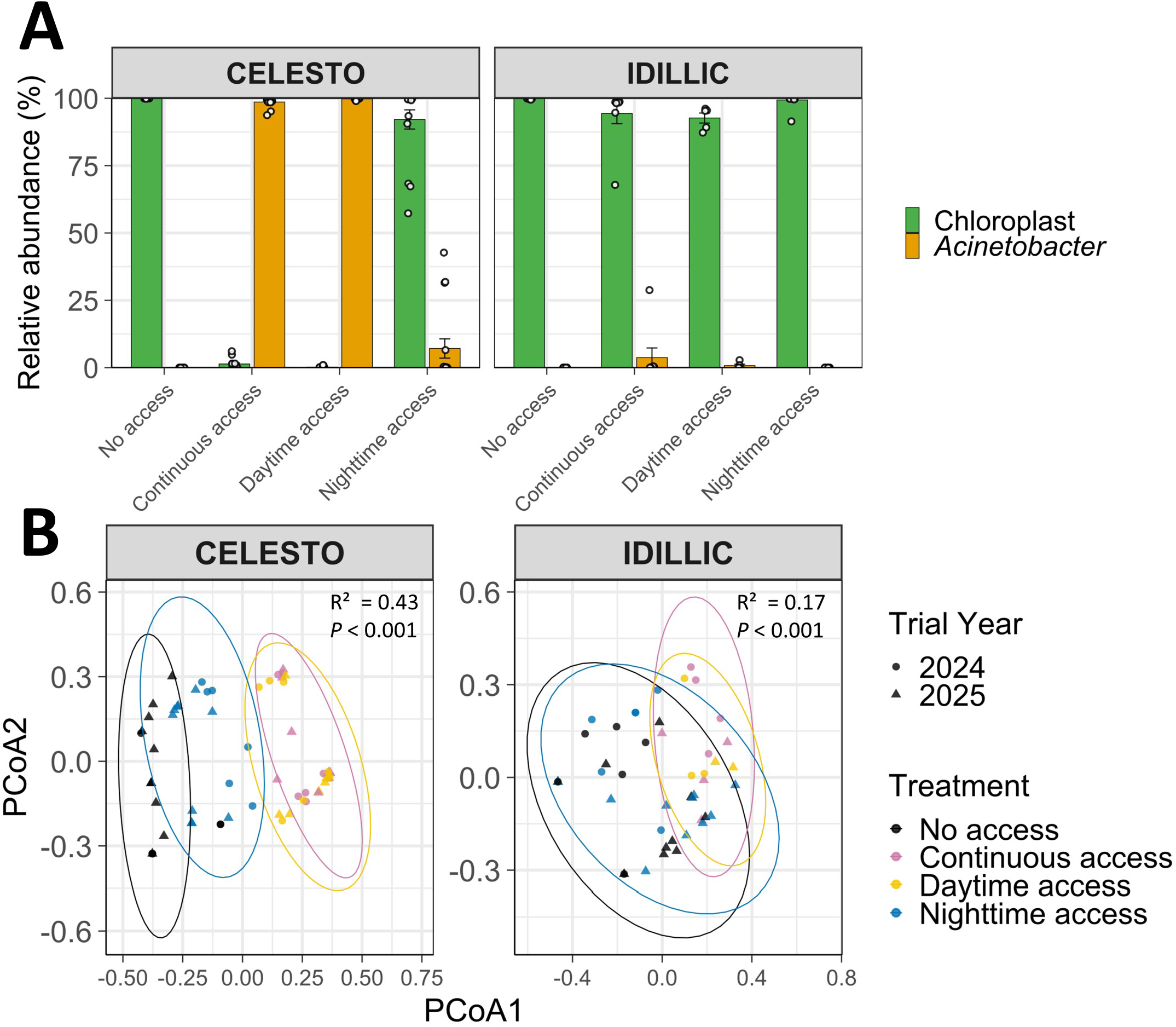
Effect of pollinator access treatment on nectar bacterial community composition in two sunflower cultivars. A. Mean relative abundance (± SE) of *Acinetobacter* and chloroplast-derived 16S sequences across pollinator access treatments (no access, continuous access, daytime access, nighttime access) in nectar samples of the Celesto and Idillic cultivars. Individual sample values are overlaid as points. Chloroplast sequences originate from co-amplification of plastidial 16S rRNA genes from residual floral tissue or pollen and do not represent bacterial taxa. n = 5–16 samples per treatment × cultivar combination. B. Principal coordinates analysis (PCoA) of bacterial community composition based on unweighted UniFrac distances, shown separately for each cultivar. Each point represents one sample, coloured by pollinator access treatment, with shape indicating trial year. Ellipses represent 95% confidence intervals. PERMANOVA marginal effects for treatment: Celesto R² = 0.43, *P* < 0.001; Idillic R² = 0.17, *P* < 0.001.

Treatment significantly influenced β-diversity in both cultivars (Figure 3B; chloroplast-derived OTUs were retained in all 16S analyses, see Methods). The effect was especially pronounced with the Celesto cultivar (PERMANOVA on unweighted UniFrac values, *P* < 0.001, R² = 0.43), with separation of the four treatments along the first PCoA axis (Figure 3B). Trial year had a significant effect in Idillic (R² = 0.08, *P* < 0.001) but not in Celesto (R² = 0.01, *P* = 0.214), suggesting that nectar communities in Idillic were more sensitive to environmental parameters.

### Pollinator visits impacted the α-diversity of nectar fungal communities differently in both cultivars

The 104 ITS samples were represented on average by 12 OTUs per sample (mean depth 22,767 reads per sample). Flower access to pollinators affected fungal α-diversity in a cultivar-dependent manner, with a significant treatment × cultivar interaction for Richness and ENS1 (ANOVA, Richness: P = 0.004; ENS1: P = 0.044), but not for ENS2 (P = 0.141).

Samples of Idillic nectar under continuous-access showed the lowest fungal diversity across all three α-diversity metrics, and differed significantly from nighttime-access (Tukey-adjusted pairwise contrast, Richness: P = 0.012; ENS1: P = 0.017; ENS2: P = 0.036) (Figure 4A). The daytime-access and no-access treatments resulted in intermediate α-diversity metric values (Figure 4A, P > 0.05 vs. nighttime-access or vs. continuous-access for either treatment across all metrics). In nectar of the Celesto cultivar, none of the pairwise contrasts between treatments reached significance for any of the three metrics (Tukey-adjusted, all P > 0.05, Figure 4A). We did not detect significant differences in any of the α-diversity metrics (Hill numbers 0, 1, and 2) between the fungal communities of no-access flowers of Celesto vs. Idillic (ANOVA, P = 0.146, 0.119 and 0.129 for q = 0, 1 and 2), ruling out a possible confounding effect of the larger nectar volume samples of Celesto on diversity measures.

**Figure 4.**
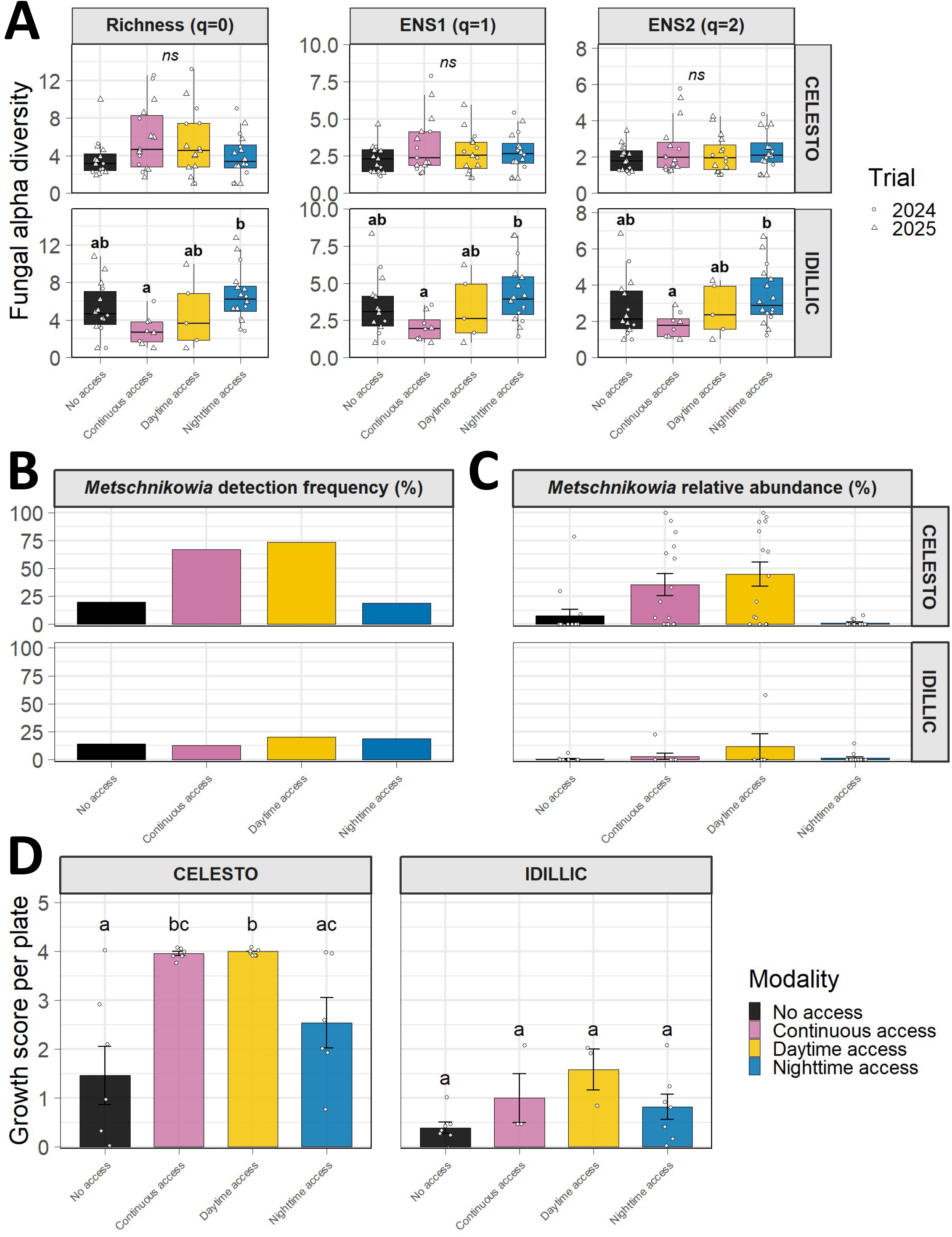
Fungal community diversity, *Metschnikowia* colonisation, and culturable microbial load in sunflower nectar across pollinator access treatments. **A.** Coverage-standardised Hill diversity metrics (q = 0, 1, 2) for Celesto and Idillic nectar fungal communities across four access treatments. Points are individual samples (circles: 2024, triangles: 2025). Letters denote significantly different groups (Šidák-adjusted pairwise contrasts within cultivar, *P* < 0.05). B, C. Detection frequency (B) and mean relative abundance (C) of *Metschnikowia* spp. across treatments in Celesto and Idillic; (C) bars show means ± SE, points show per-sample values. D. Culturable microbial load (mean growth score across four media — TSA, R2A, PDA, YM; 0–4 scale) per sample for Celesto and Idillic across treatments. Letters denote significantly different groups (pairwise Wilcoxon rank-sum tests on per-sample means within cultivar, Benjamini–Hochberg corrected, *P*.adj < 0.05).

### Pollinator visits impacted nectar fungal β-diversity in a cultivar-dependent manner

Nectar fungal community composition of the Celesto cultivar differed significantly between treatments for three of the four β-diversity metrics (PERMANOVA, Bray–Curtis, unweighted UniFrac and weighted UniFrac: all P < 0.001; Jaccard: P = 0.114; Bray–Curtis R² = 0.088, Jaccard R² = 0.056, unweighted UniFrac R² = 0.088, weighted UniFrac R² = 0.137; Figure S4). Trial year had a significant effect on Jaccard and unweighted UniFrac (both P ≤ 0.004), and to a lesser extent on Bray-Curtis (P = 0.020), but not on weighted UniFrac (P = 0.156). Daytime- and continuous-access samples did not differ from each other (all *P* > 0.49). Both daytime- and continuous-access differed significantly from the no-access and nighttime-access treatments on three of the four metrics (all but Jaccard, q < 0.027). Relative to no-access, nighttime access affected the two abundance-weighted metrics (Bray-Curtis, weighted UniFrac: q < 0.044) but not the two presence/absence metrics (Jaccard, unweighted UniFrac: q > 0.37). In contrast, pollinator access treatment had no effect on fungal community composition in nectar of the Idillic flowers (*P* > 0.05 for all β-diversity metrics), but trial year significantly structured composition (*P* ≤ 0.001 for all metrics, R² = 0.041–0.106). No pairwise comparison remained significant in Idillic after correction. Together, these data indicate that the β-diversity of fungal communities followed similar trends to bacterial communities, with communities of the Celesto flowers being more responsive to treatment than Idillic communities, and daytime exposure specifically driving this compositional shift. The significant effect of trial year indicates that nectar fungal communities were more sensitive to the environment than bacterial communities for both cultivars.

In particular, the combined abundance of *Metschnikowia* OTUs differed markedly between cultivars and treatments (Figure 4B). We detected *Metschnikowia* OTUs in nectar samples of the Idillic cultivar at frequencies between 12.5% and 20.0% across samples, with a mean relative abundance between 0.5% and 11.6% of the total ITS reads. Fungal communities of the Celesto cultivar, however, were dominated by *Metschnikowia*, with OTUs found in 73.3% of daytime-access samples (mean relative abundance = 44.8%), 66.7% of continuous-access samples (35.5%), 18.8% of nighttime-access samples (1.1%) and 20.0% of no-access samples (7.8%; family-level composition shown in Figure S5).

Together, these data suggest that the nectars of Idillic and Celesto offer distinct microbial niches, with the Celesto cultivar favouring specialist yeasts and bacteria associated with diurnal visits. In contrast, nectar of the Idillic cultivar appears to favour less predictable microbial assemblages upon pollinator visitation regimes.

### Pollinator visits affected culturable microbial loads differently in both cultivars

The contrasted nectar communities according to pollinator exclusion conditions might also be explained by a different capacity of nectar to sustain the growth of microbes inoculated by diurnal pollinators. To test this, we spread nectar samples collected using the same protocol as for microbiota sequencing, but from a separate set of plants from both cultivars on generalist bacteriological (TSA, R2A) and mycological (PDA, YM) culture media (Table S5). Samples collected from Celesto flowers under both continuous- and daytime-access regimes reached near-maximal growth scores (mean 3.96 ± 0.10 and 4.00 ± 0.00 on the 0–4 scale), significantly higher than under no-access (mean 1.46 ± 1.58) (n = 6–7 per treatment; Wilcoxon rank-sum test, BH-adjusted, q < 0.05, Figure 4C). Nighttime-access samples had intermediate microbial loads. In contrast, samples collected from Idillic flowers always yielded few colonies, regardless of treatment (Figure 4C). Together, these data suggest that the nectar of Idillic flowers is overall less permissive to the growth of microbes brought by pollinator visits.

### Visit frequency alone did not predict community structure

Alternatively, we tested whether the frequency of visits was associated with nectar community composition for each cultivar. Neither the number of bee and bumblebee visits nor moth visit rates correlated with α-diversity measures in either cultivar (Idillic diurnal, n = 12, *P* = 0.450–0.549; Celesto nocturnal, n = 27, *P* = 0.182–0.302; Idillic nocturnal, n = 21, *P* = 0.558–0.972; Figure S6). Celesto diurnal visits showed a negative trend with ENS1 and ENS2 that did not reach significance (*P* = 0.187 and *P* = 0.214, respectively; Richness *P* = 0.148). Overall, visit frequency alone was not sufficient to predict nectar fungal diversity. Beta-diversity showed the same pattern: visit rate did not predict community composition in either cultivar or time window across all 4 dissimilarity metrics (PERMANOVA, all P ≥ 0.074).

### Access treatment shifted individual floral volatiles without reshaping overall blend composition

To test whether floret VOC composition is affected by cultivar and/or accessibility of pollinators, we analysed the VOC profiles of differently treated florets of both cultivars (dataset available at https://doi.org/10.57745/LGKSOT). Multivariate principal coordinates analysis (PCoA) on Bray-Curtis distances of 111 detected VOCs from 54 plants revealed clear separation of samples according to genotype along PCo1 (39% of the variance explained, Figure 5). This pattern was corroborated by PERMANOVA on Bray–Curtis dissimilarities, which detected a significant effect of cultivar (R² = 34.7%, *P* < 0.0001, betadisper/permutest, *P* = 0.328). VOCs that were more abundant in Celesto compared to Idillic seemed to belong to the monoterpenoids class, while Idillic was characterized by VOCs having a sesquiterpenoid signature. However, access treatment had no detectable effect (PERMANOVA *P* = 0.14 for Celesto; *P* = 0.11 for Idillic).

**Figure 5.**
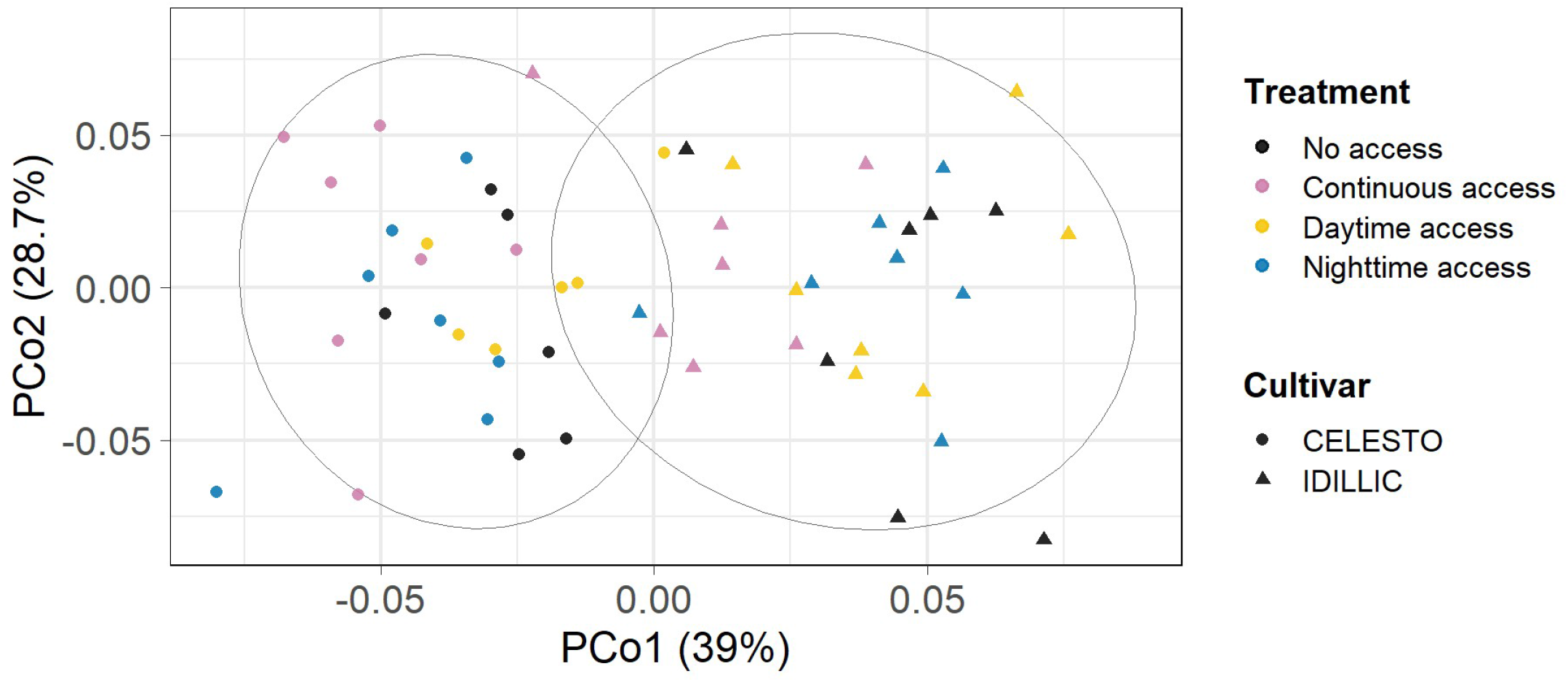
Floral volatile blend composition across cultivars and access treatments. Principal coordinates analysis (PCoA) of floral volatile blend composition (Bray–Curtis dissimilarity on relative abundances of the 111 compounds detected) across all floret samples. Shape indicates cultivar; colour indicates pollinator access. Ellipses represent 95% confidence intervals around each cultivar’s samples. Axis labels indicate the percentage of total variance explained by each principal coordinate.

In the absence of a global effect of treatment on volatile profiles, we asked whether individual compounds differed between access treatments within each cultivar, focusing on the no-access vs. continuous-access contrast, which captured most of the significant compound-level differences (Table S8). Four compounds in Celesto and 8 of the 10 significant compounds in Idillic were significantly reduced under continuous access relative to no access (q < 0.05; Figure S7). Of the compounds reduced under continuous access, two were consistent with a microbial origin — α-guaiene (feature 6262, significant in Celesto) and a related compound (feature 6553, significant in Idillic).

Overall, floral volatile blends of florets were characterised by mono- and sesquiterpenoid compounds. The VOC composition differed markedly between the two cultivars, whereas pollinator-access treatment had only minor effects.

## DISCUSSION

Netting flowers by day or by night partitioned the visitor guilds effectively, with bees and bumblebees almost absent from nighttime-access plants, and moths from daytime-access plants, in both cultivars. Camera monitoring confirmed that both cultivars received visits from the same pollinator guilds under each access regime. Separating the guilds showed that they did not contribute equally to nectar colonisation, and that this asymmetry depended on the cultivar receiving the visits. In Celesto, pollinator access reshaped the overall composition of the fungal community, while diversity itself remained comparable across treatments. In Idillic, the reverse held: continuous access lowered fungal diversity relative to nighttime access, while fungal community composition was not detectably affected by treatment. Composition was, however, structured by trial year, suggesting Idillic nectar communities responded more to environmental variation than to pollinator guild identity. This was not simply a matter of how much visitation occurred: photographic monitoring showed that visit frequency, whether diurnal or nocturnal, did not predict fungal diversity or community composition in either cultivar. Floral traits therefore appear to gate the outcome of pollinator visits as much as the pollinators themselves.

Where community composition shifted, it was driven by specialist taxa reaching dominance under diurnal-access regimes. In Celesto, both the yeast *Metschnikowia* and the bacterium *Acinetobacter* dominated under continuous- and daytime-access, while remaining present but non-dominant (*Metschnikowia*) or nearly absent (*Acinetobacter*) under no-access and nighttime access. This establishment tracked the diurnal guild specifically, rather than just insect exposure. This is consistent with bees being the principal vectors of both *Metschnikowia* and nectar-specialist *Acinetobacter* in wild systems [6,9,11,34]. In addition, *Metschnikowia* and *Acinetobacter* are similarly reported to co-occur in nectar [35]. Niche partitioning (e.g. preferential use of glucose vs fructose), as previously hypothesized [36], would explain the co-occurrence of both bacteria and yeast taxa. In Idillic, by contrast, neither taxon reached dominance under any treatment: *Metschnikowia* OTUs were detected in only a fifth of daytime-access flowers and remained at low relative abundance when present, while *Acinetobacter* OTUs remained rare. This absence of dominance was mirrored in culturable load: Celesto nectar reached near-maximal growth scores under daytime and continuous access, while Idillic nectar yielded few colonies across all treatments regardless of exposure. The two access windows also differed in duration (16 h daytime vs 8 h night) and temperature, so part of the day/night asymmetry may reflect growth opportunity rather than inoculum composition alone.

What allowed these specialists to establish in Celesto but not Idillic may lie in the nectar itself. Idillic florets held roughly four times less nectar than Celesto and a correspondingly smaller sugar mass. Its nectar also retained a substantial sucrose fraction, where Celesto nectar was almost entirely hexose-based, a difference that may also bear on which taxa can establish. Nectar-specialist yeasts and bacteria grow in a volume that visitors themselves deplete, so a floret producing less nectar offers a smaller resource to colonise. In *Epilobium*, plants with higher nectar volumes were more likely to contain bacteria [16], consistent with what we observed for *Acinetobacter*. Furthermore, nectar of bee-pollinated species is a flow-through resource: it is secreted over the lifetime of a floret, removed by visitors, and re-secreted after removal [37,38]. The two cultivars thus differed not only in standing volume but, in combination with their contrasting visitation rates, in the flux of nectar through the floret. Taking the product of mean floret volume and mean diurnal visit rate as a proxy, and assuming that all nectar is removed after each bee visit, that flux is roughly an order of magnitude higher in Celesto (∼13-fold). Higher numbers of visits increase the number of inoculation events, but the number of propagules brought by each event is likely modest, fewer than a hundred yeast cells [39] and unlikely to compensate for the depletion of established communities. These repeated cycles of nectar secretion and depletion are conceptually similar to cycles of dilution followed by growth: both theory and co-culture experiments show that fast-growing organisms are favoured under low-density, high-dilution conditions [40]. This is consistent with the higher flux, and thus higher effective dilution rate, experienced by Celesto nectar communities. Although the influence of secretion/depletion cycles on realized growth has been modelled for *Metschnikowia*, their impact on competition outcomes within nectar communities remains an open question.

While we observed sharp differences in the response of nectar microbial communities to insect visits in Celesto, no such strong effect on the overall floral VOC profiles was found. Instead, cultivar was the strongest driver of blend composition. In Celesto, linear models identified 4 VOCs with a significant treatment effect, all showing reduced abundance under continuous access relative to no access. A similar pattern was observed for 8 of the 10 significant VOCs in Idillic. Although this could be attributed to netting-induced differences in flower microclimate, it is also consistent with post-pollination declines in floral scent reported in other systems. This decline could serve as an olfactory cue that enables foragers to discriminate depleted from unvisited flowers, as shown by lower bee visitation to pollinated than to unpollinated thistle flower heads [41].

Following a similar trend, we observed an apparent dilution of the microbial-associated α- guaiene signal under continuous vs no-access in both cultivars. Volatile α-guaiene has previously been associated with yeast fermentation in other systems [42]. This is also consistent with a model of repeated secretion/depletion cycles during pollination windows, effectively leaving less time for microbial signatures to accumulate in open flowers, particularly in Celesto, where nectar flux is highest.

In conclusion, this study shows that pollinator guilds shape nectar microbiome assembly differently depending on cultivar, reshaping community composition in Celesto, shifting diversity without restructuring composition in Idillic. This guild- and context-dependent restructuring did not, however, extend significant shifts in flower VOC profiles. Deciphering the mechanisms that drive these differences will require connecting the physiological aspects of nectar chemistry/secretion, pollinator biology and microbial ecology.

## ACKNOWLEDGMENTS

We thank Baptiste Mayjonade for his help and advice on ONT Nanopore sequencing, Mélanie Carcagno for her assistance with nectar bacterial isolation, Nicolas Pouilly for the floret grinding protocol, and the CNRGV for the free loan of the grinding jars. Insect silhouettes in Figure 4B are from PhyloPic (phylopic.org): *Apis mellifera and Bombus terrestris* by Melissa Broussard (CC BY 4.0 and CC BY 3.0, respectively), and *Autographa gamma* by Gareth Monger (CC BY 3.0); modified (recoloured, resized). We also thank Henriëtte van Eekelen, Bert Schipper and Dirk Bosch for their support with the floret VOC analyses.

## FUNDING

This work is part of HELEX project funded by the European Union’s Horizon Europe Research and Innovation Actions programme under grant agreement N°101081974. This research used the PHENOME-EMPHASIS facility Phenotoul-Polliphen (Phenome-ANR-11-INBS-0012, https://doi.org/10.15454/1.5483266728434124E12) and was part of the French Laboratory of Excellence project “TULIP” (ANR-10-LABX-41; ANR-11-IDEX-0002-02).

This work was funded by the Plant2Pro Carnot Institute in the frame of its 2022 call for projects. Plant2Pro is supported by ANR (agreement #22-CARN-024-01 – 2021). G.T. acknowledges support from the INRAE BAP department.

## CONTRIBUTIONS

G.T., N.L., and A.C. conceived the study. G.T., O.R. managed field trial, sampled nectar and florets, and deployed the cameras. M.P-J quantified nectar traits. G.T. and O.R. isolated nectar samples on culture media and scored colony growth. R.M., and K.C. developed the floret collection protocol, performed the volatile analysis and identified the compounds. G.T., O.R. extracted nucleic acids from nectar samples, performed PCR and sequencing. G.T. and A.C. analysed amplicon sequencing data. G.T. evaluated the detection model. G.T. performed statistical analyses. G.T., A.C., and N.L. wrote the manuscript with contributions from all authors. A.C. and N.L. contributed equally to the supervision of the work.

## DATA STATEMENT

All sequence data generated during the study are available in a public repository, accession numbers are provided in the main text.

Raw data on nectar traits (volume, sugar composition), pollinator visitation counts, microbial culturability scores and associated plate photographs, and the reference sequences underlying *Metschnikowia* OTU merging are available on the INRAE institutional repository at https://doi.org/10.57745/LGKSOT.

Analysis code is available at https://github.com/GuillaumeTueux/sunflower-nectar-pollinator-exclusion.

## COMPETING INTERESTS

The authors declare no competing interests.

## DECLARATION OF GENERATIVE AI AND AI-ASSISTED TECHNOLOGIES IN THE MANUSCRIPT PREPARATION PROCESS

During the preparation of this work the author(s) used Anthropic Claude Sonnet 5 in order to draft R code used for some figures, and for proofreading of the manuscript. After using this tool/service, the author(s) reviewed and edited the content as needed and take(s) full responsibility for the content of the published article.

## Notes

https://doi.org/10.57745/LGKSOT

### Competing Interest Statement

The authors have declared no competing interest.

