## Supplementary material for "Same visitors, different outcomes: floral phenotype gates pollinator-vectored nectar microbial establishment": Figure S

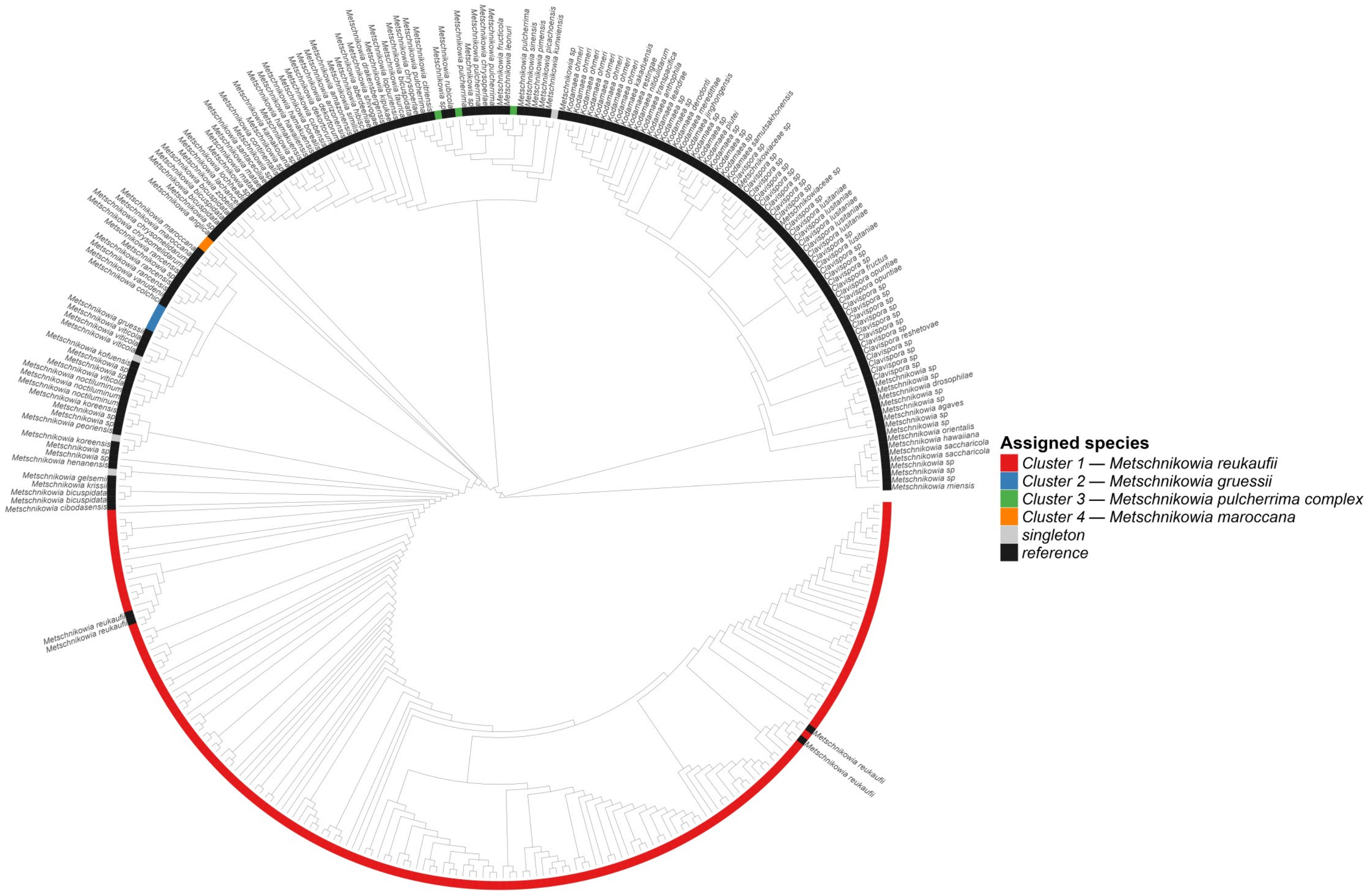


**Figure S1. Maximum likelihood phylogeny of Metschnikowia OTUs and reference sequences used for phylogeny-informed merging.** The phylogeny was inferred from 184 OTU sequences and 169 ITS reference sequences of Metschnikowia and related genera (Clavispora, Kodamaea), aligned with MAFFT v7.505 and trimmed with Trimal, using FastTree2 (GTR+GAMMA model) and rooted at the midpoint. The tree is displayed as a cladogram (branch lengths not proportional to substitutions). The outer ring indicates cluster assignment: OTUs forming clades with maximum pairwise patristic distance below the merging threshold were merged into 4 clusters (colours) or retained as singletons (grey), starting with the largest candidate clades. Reference sequences are shown in black. Each cluster is annotated with the nearest reference species, defined as the species present within the minimal clade containing all its OTUs or, when absent, showing the minimum mean patristic distance across all OTU members. Tip labels show species names for reference sequences only. For cluster 3, reference sequences from seven nominal species of the Metschnikowia pulcherrima complex fall within the same clade, precluding assignment to a single species.

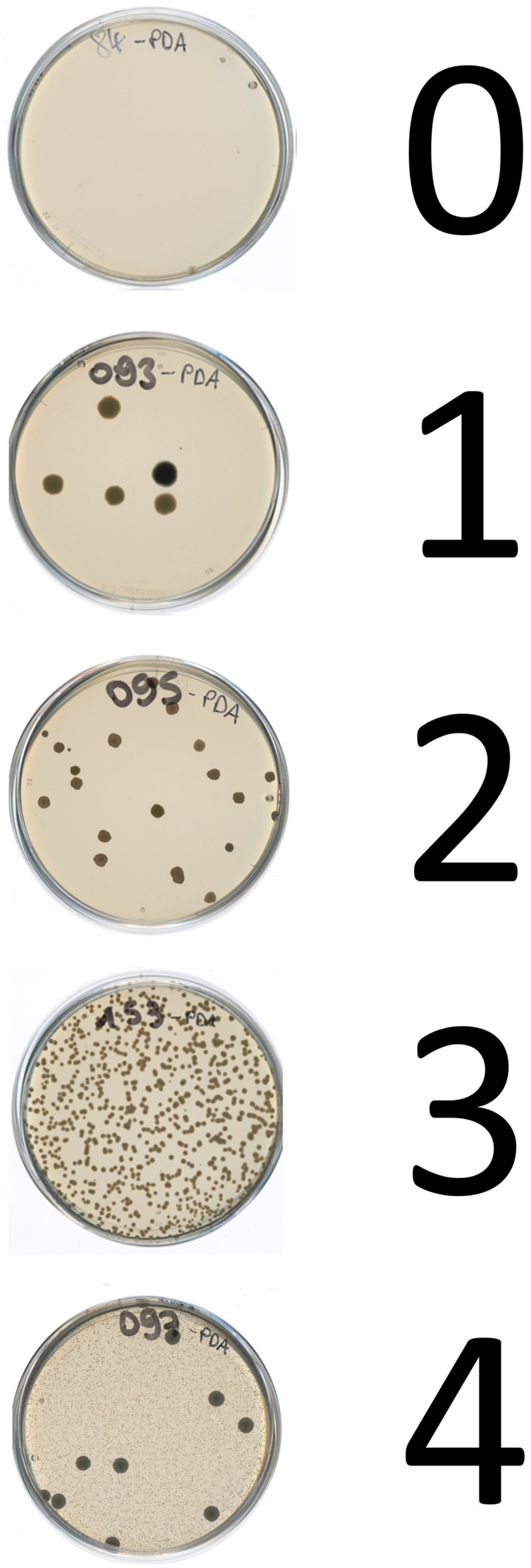


**Figure S2. Reference photographs illustrating the semi-quantitative growth scoring scale used for nectar culture plates (potato dextrose agar, PDA).** Scores: 0, no colonies; 1, 1–10 colonies; 2, 11–50 colonies; 3, 51–500 colonies; 4, >500 colonies (confluent lawn). Scale applies to all four culture media used in this study (TSA, R2A, PDA, YM agar).

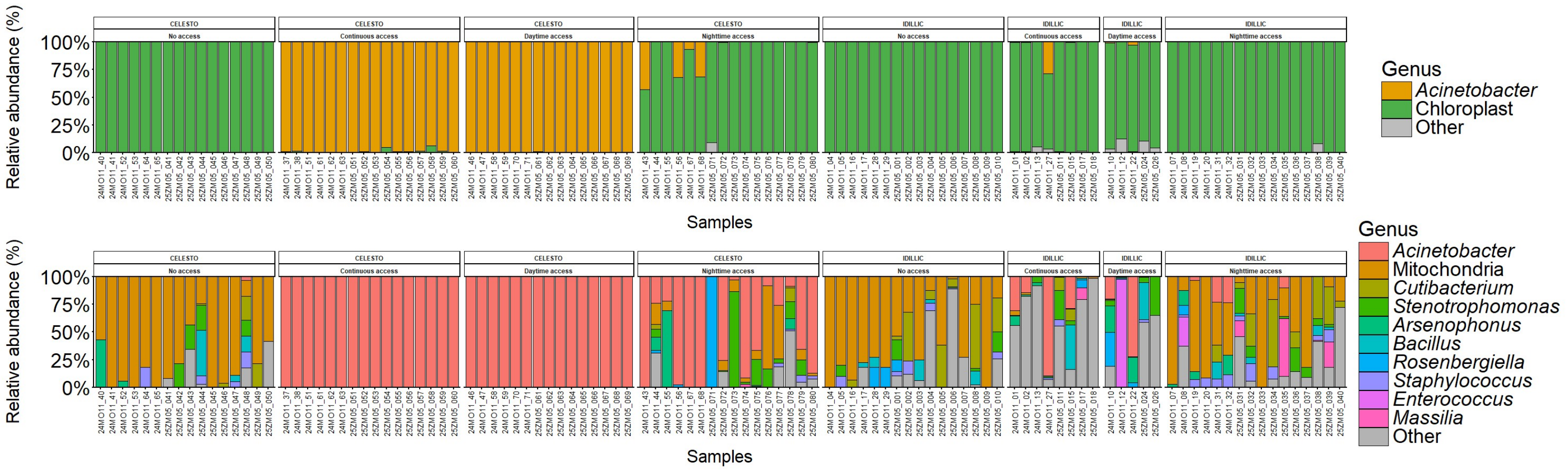


**Figure S3. Relative abundance of bacterial genera across all retained 16S samples (n = 109), facetted by cultivar (CELESTO, IDILLIC) and pollinator access treatment (No access, Continuous access, Daytime access, Nighttime access).** Each bar represents one sample. Top : Only *Acinetobacter* and chloroplast-derived sequences are shown individually, all remaining genera are pooled as "Other". Bottom : Genus-level composition after subtraction of chloroplast-derived sequences, the ten most abundant genera are shown individually, all remaining genera pooled as "Other".

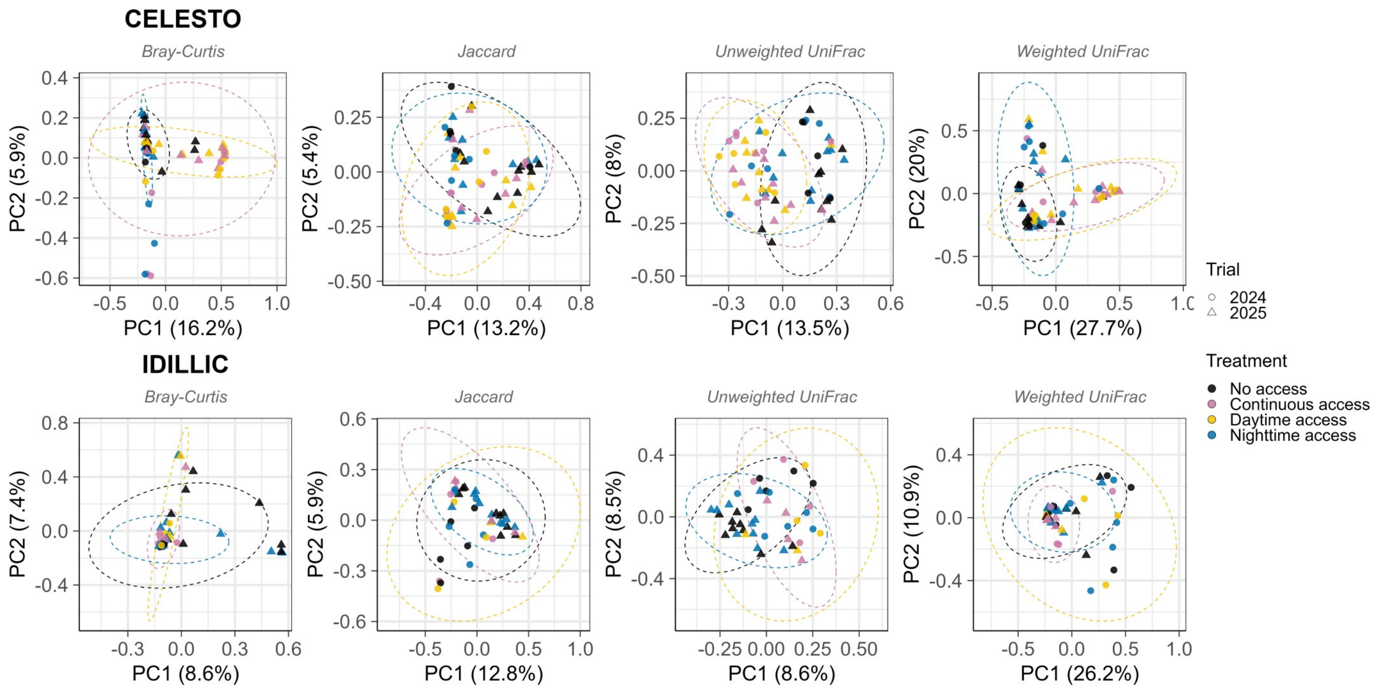


**Figure S4. Principal coordinates analysis (PCoA) of fungal nectar community composition in CELESTO (top row) and IDILLIC (bottom row) under four pollinator access treatments, across four beta-diversity metrics: Bray-Curtis dissimilarity (abundance-based), Jaccard dissimilarity (presence/absence), unweighted UniFrac (phylogenetic, unweighted), and weighted UniFrac (phylogenetic, abundance-weighted).** Points represent individual samples. Shape indicates trial year. Colour indicates pollinator access treatment. Dashed ellipses show 95% confidence regions per treatment.

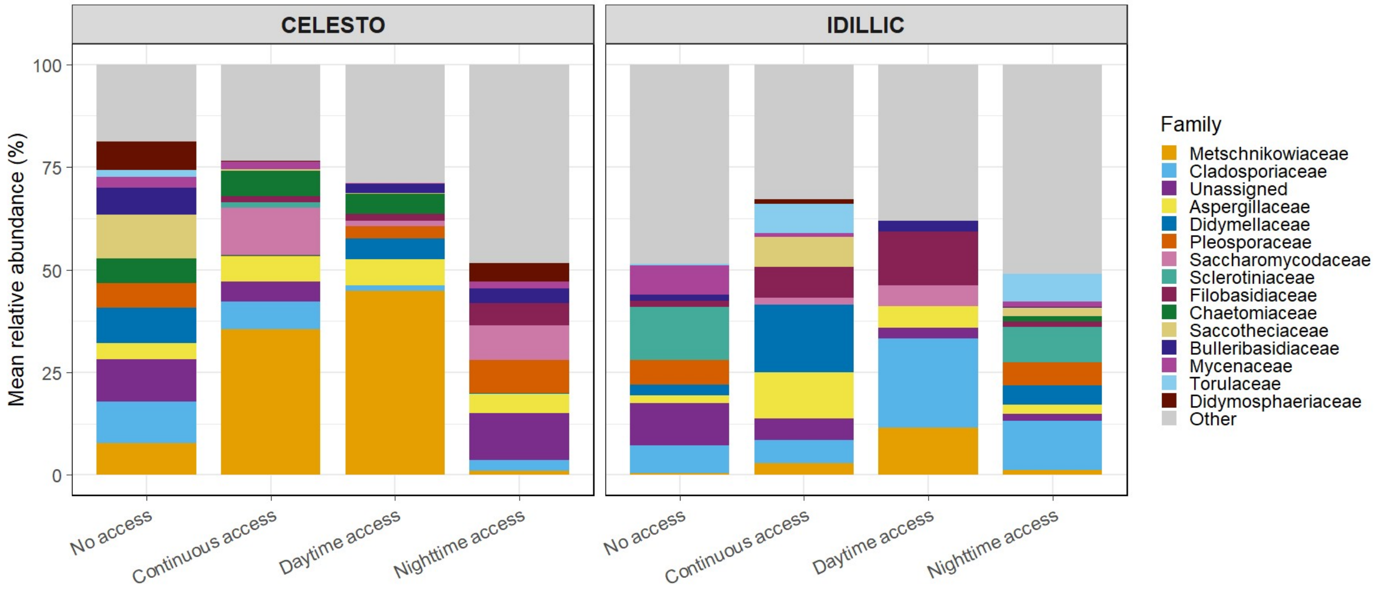


**Figure S5. Fungal community composition at the family level across visitor treatments and cultivars.** Stacked bar plots show mean relative abundance (%) of the 15 most abundant fungal families across all samples, plus all remaining families pooled as "Other" (grey). Values represent means across samples within each visitor treatment × cultivar combination. Colours identify fungal families as indicated in the legend.

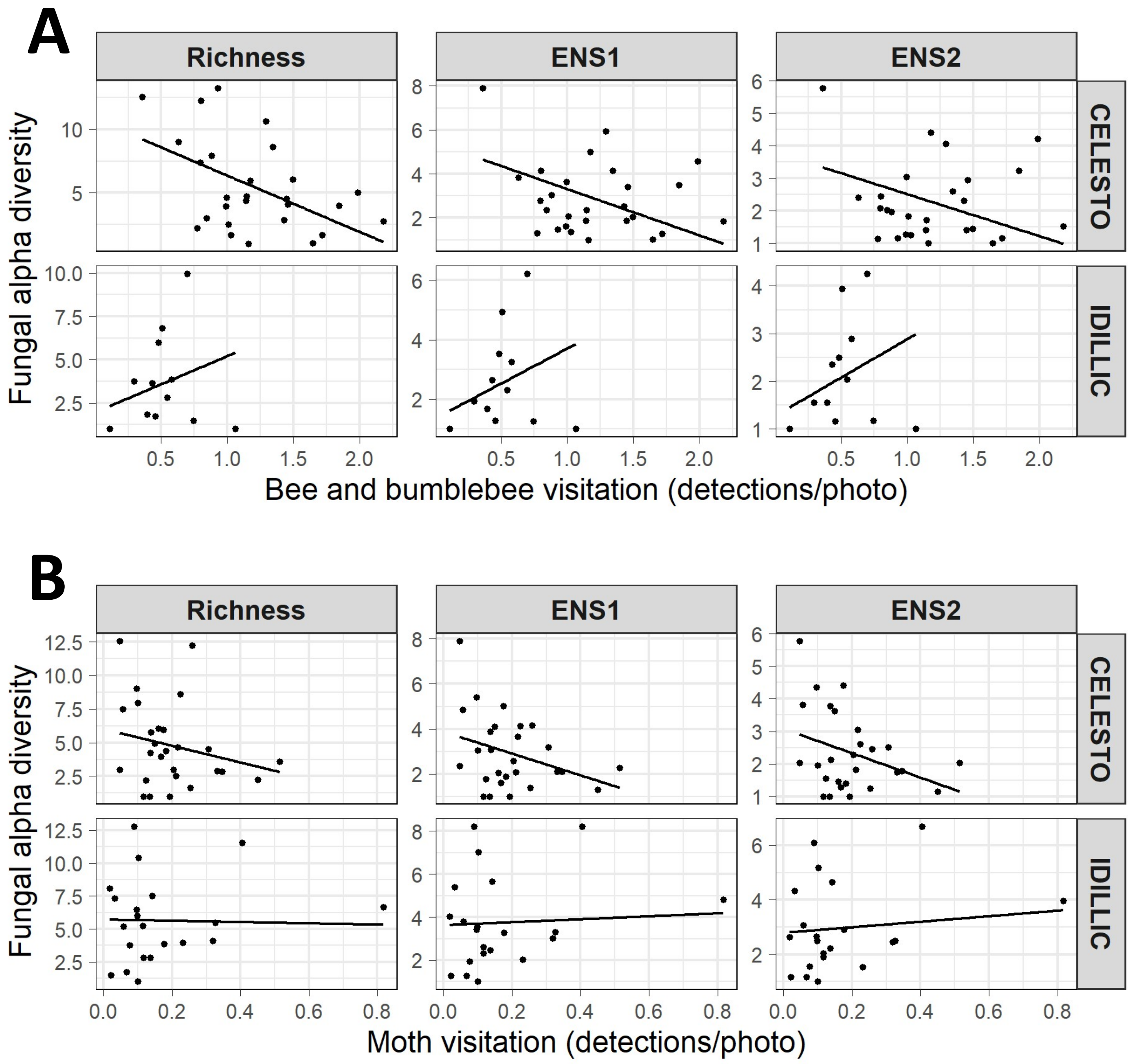


**Figure S6. Relationship between pollinator visitation and fungal alpha diversity (Richness, ENS1, ENS2) in CELESTO and IDILLIC.** (A) Diurnal visitation (bee and bumblebee detections/photo), restricted to daytime and continuous access treatments. (B) Nocturnal visitation (moth detections/photo), restricted to nighttime and continuous access treatments. Points show per-sample raw values. Lines show model-adjusted predictions averaged over the observed distribution of treatment and trial year.

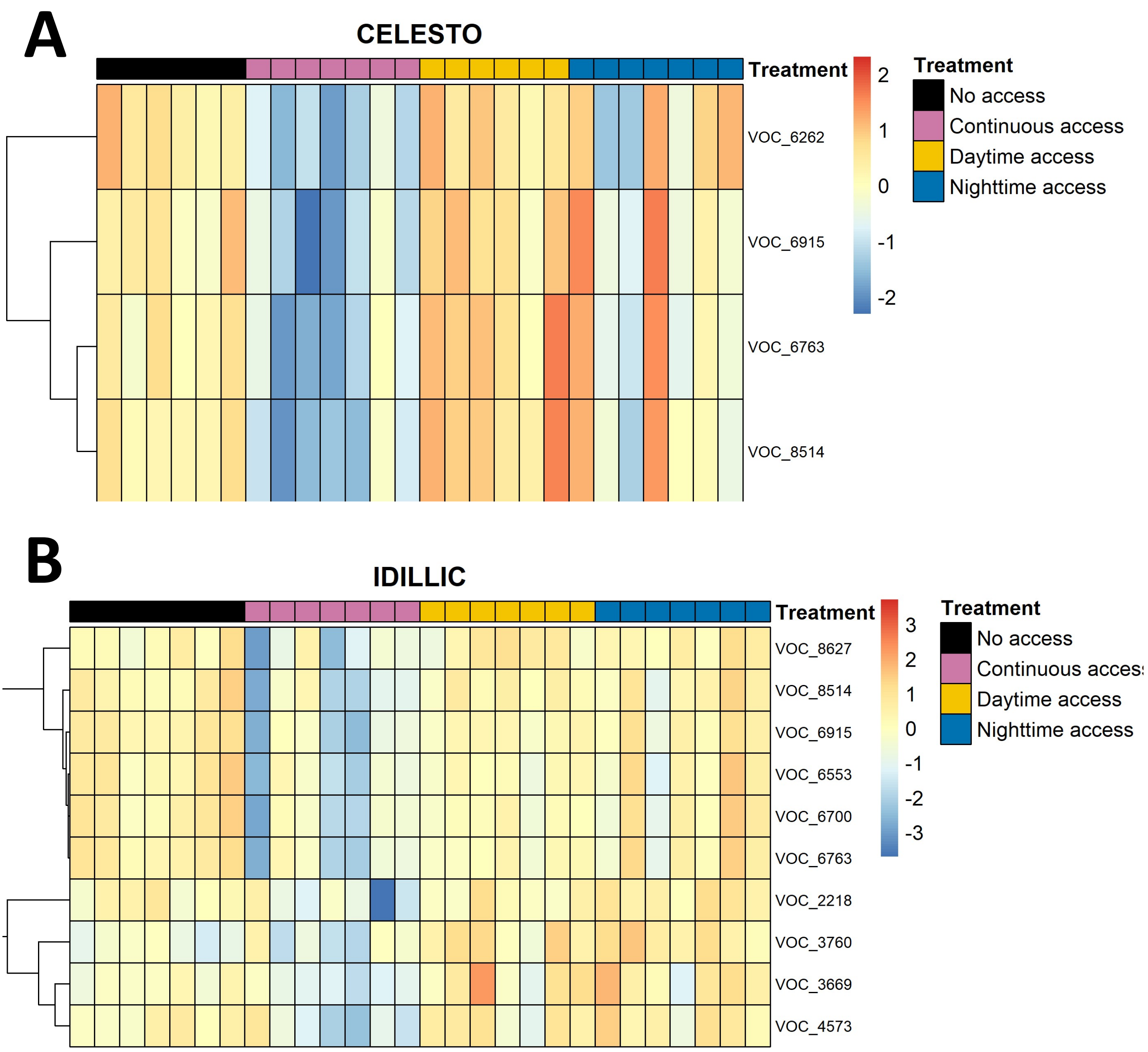


**Figure S7. Univariate compound-level responses to pollinator access treatment, by cultivar.** Heatmap of significant volatile compounds in (A) Celesto and (B) Idillic. Row-scaled (z-score) log10(abundance + 1) values for volatile compounds showing a significant effect of pollinator access modality (ANOVA, Benjamini–Hochberg-adjusted model $q < 0.05$) within each cultivar (Celesto, n = 4 compounds; Idillic, n = 10 compounds; see Table S8 for Šidák-adjusted pairwise groupings). Row scaling was computed independently within each cultivar. Columns represent individual samples, ordered by pollinator access treatment within each cultivar. Rows are hierarchically clustered by correlation distance, independently within each cultivar. Column annotation indicates pollinator access treatment.
