## Supplementary material for "Same visitors, different outcomes: floral phenotype gates pollinator-vectored nectar microbial establishment": Table S

| Class | True Positive | False Positive | False Negative | Precision | Recall | F1 |
|---|---|---|---|---|---|---|
| Bee | 2480 | 19 | 343 | 99.24% | 87.85% | 93.20% |
| Bumblebee | 174 | 3 | 22 | 98.31% | 88.78% | 93.30% |
| Moth | 301 | 64 | 54 | 82.47% | 84.79% | 83.61% |

**Table S1. Box-level validation of pollinator detection and classification.** Per-class true positive (TP), false positive (FP) and false negative (FN) bounding-box counts for the YOLO11x-based detection model (Pollicrop), assessed against expert manual annotation on a validation subset of 5,000 images. Precision, recContinuous and F1 score were calculated as TP/(TP+FP), TP/(TP+FN) and 2TP/(2TP+FP+FN), respectively, per class.

| Culti-var | Treatment | Trial year | n plants microbiota | n plants camera | Total photos | Total bee | Total bumblebee | Total moth | Total photos Day | Total photo Night |
|---|---|---|---|---|---|---|---|---|---|---|
| CELES-TO | Continuous access | 2024 | 6 | 6 | 17032 | 8192 | 278 | 1168 | 10820 | 6212 |
| CELES-TO | Continuous access | 2025 | 10 | 8 | 20961 | 14631 | 359 | 1634 | 12711 | 8250 |
| CELES-TO | Daytime access | 2024 | 6 | 5 | 8836 | 7980 | 261 | 23 | Not applicable | Not applicable |
| CELES-TO | Daytime access | 2025 | 10 | 10 | 14702 | 22935 | 1312 | 48 | Not applicable | Not applicable |
| CELES-TO | Nighttime access | 2024 | 6 | 5 | 5206 | 2 | 24 | 716 | Not applicable | Not applicable |
| CELES-TO | Nighttime access | 2025 | 10 | 9 | 9308 | 145 | 73 | 2224 | Not applicable | Not applicable |
| CELES-TO | No access | 2024 | 6 | 0 | 0 | 0 | 0 | 0 | Not applicable | Not applicable |
| CELES-TO | No access | 2025 | 10 | 0 | 0 | 0 | 0 | 0 | Not applicable | Not applicable |
| IDILLIC | Continuous access | 2024 | 4 | 4 | 11450 | 2865 | 377 | 405 | 7334 | 4116 |
| IDILLIC | Continuous access | 2025 | 4 | 4 | 9935 | 4671 | 909 | 326 | 6234 | 3701 |
| IDILLIC | Daytime access | 2024 | 3 | 3 | 5342 | 2087 | 287 | 51 | Not applicable | Not applicable |
| IDILLIC | Daytime access | 2025 | 2 | 2 | 3054 | 1099 | 125 | 0 | Not applicable | Not applicable |
| IDILLIC | Nighttime access | 2024 | 6 | 5 | 4906 | 1 | 0 | 959 | Not applicable | Not applicable |
| IDILLIC | Nighttime access | 2025 | 10 | 9 | 9148 | 15 | 64 | 1919 | Not applicable | Not applicable |
| IDILLIC | No access | 2024 | 6 | 0 | 0 | 0 | 0 | 0 | Not applicable | Not applicable |
| IDILLIC | No access | 2025 | 10 | 0 | 0 | 0 | 0 | 0 | Not applicable | Not applicable |

**Table S2. Sample sizes for nectar microbiota sequencing and camera-trap pollinator visitation, by cultivar, pollinator access treatment and trial year.** n plants microbiota indicates the number of plants sampled for nectar microbiota sequencing, prior to marker-specific quality filtering. n plants camera indicates the number of plants with usable camera-trap image data, following exclusion of plants affected by camera malfunction (flash failure, incorrect date stamp, or misalignment). Total photos, total bee, total bumblebee and total moth report the summed number of images and pollinator detections across Continuous cameras within each cultivar × treatment × trial year combination. Total photos day and total photos night report the number of images fContinuousing within the daytime (start of monitoring to 22:00) and nighttime (22:00 to end of monitoring) windows respectively, applicable only to the continuous access treatment. No access plants had no camera deployed, hence zero values across camera-derived columns.

| Primers | |
|---|---|
| 16S-tailed_F : | TTTCTGTTGGTGCTGATATTGCAGRGTTYGATYMTGGCTCAG |
| 16S-tailed_R : | ACTTGCCTGTCGCTCTATCTTCRGYTACCTTGTTACGACTT |
| ITS4-tailed_R : | ACTTGCCTGTCGCTCTATCTTCTCCTCCGCTTATTGATATGC |
| ITS1F-tailed_F : | TTTCTGTTGGTGCTGATATTGCTTGGTCATTTAGAGGAAGTAA |

**Table S3. Adapter-tailed primer sequences used for long-amplicon nanopore sequencing.** Adapter-tailed primers were used to amplify the bacterial 16S rRNA gene (16S-tailed_F/16S-tailed_R) and the fungal ITS1–ITS4 region (ITS1F-tailed_F/ITS4-tailed_R). Sequences are reported 5′→3′ and include the 5′ adapter tails required for Oxford Nanopore ligation library preparation, followed by the locus-specific primer sequence.

| Amplification | Cultivar | Distance | Treatment $R^2$ | Treatment F | Treatment *P* | Treatment betadisper *P* | Trial year $R^2$ | Trial year F | Trial year *P* |
|---|---|---|---|---|---|---|---|---|---|
| 16S | IDILLIC | Bray-Curtis | 0.150 | 2.47 | 0.036 | 0.022 | 0.028 | 1.37 | 0.193 |
| 16S | IDILLIC | Jaccard | 0.141 | 2.38 | <0.001 | 0.020 | 0.063 | 3.18 | <0.001 |
| 16S | IDILLIC | Unweighted UniFrac | 0.174 | 3.10 | <0.001 | 0.733 | 0.084 | 4.51 | <0.001 |
| 16S | IDILLIC | Weighted Uni-Frac | 0.200 | 3.50 | 0.004 | 0.021 | 0.023 | 1.18 | 0.281 |
| 16S | CELESTO | Bray-Curtis | 0.977 | 975 | <0.001 | 0.004 | 0.003 | 7.89 | 0.008 |
| 16S | CELESTO | Jaccard | 0.495 | 20.4 | <0.001 | <0.001 | 0.034 | 4.14 | 0.006 |
| 16S | CELESTO | Unweighted UniFrac | 0.429 | 14.9 | <0.001 | 0.188 | 0.013 | 1.38 | 0.214 |
| 16S | CELESTO | Weighted Uni-Frac | 0.977 | 1013 | <0.001 | 0.003 | 0.002 | 7.70 | 0.008 |
| ITS | IDILLIC | Bray-Curtis | 0.067 | 0.949 | 0.698 | 0.058 | 0.041 | 1.741 | <0.001 |
| ITS | IDILLIC | Jaccard | 0.061 | 0.889 | 0.866 | 0.596 | 0.064 | 2.782 | <0.001 |
| ITS | IDILLIC | Unweighted UniFrac | 0.075 | 1.100 | 0.190 | 0.206 | 0.055 | 2.415 | <0.001 |
| ITS | IDILLIC | Weighted Uni-Frac | 0.061 | 0.935 | 0.553 | 0.674 | 0.106 | 4.813 | <0.001 |
| ITS | CELESTO | Bray-Curtis | 0.088 | 1.861 | <0.001 | 0.234 | 0.027 | 1.730 | 0.020 |
| ITS | CELESTO | Jaccard | 0.056 | 1.147 | 0.114 | 0.352 | 0.035 | 2.168 | 0.001 |
| ITS | CELESTO | Unweighted UniFrac | 0.088 | 1.863 | <0.001 | 0.247 | 0.032 | 2.013 | 0.003 |
| ITS | CELESTO | Weighted Uni-Frac | 0.137 | 3.038 | <0.001 | 0.095 | 0.022 | 1.452 | 0.156 |

**Table S4. PERMANOVA and multivariate dispersion test results for bacterial (16S) and fungal (ITS) community composition.** Analyses were conducted separately for each amplicon and cultivar using four dissimilarity metrics. PERMANOVA was performed with 99,999 permutations using marginal tests (adonis2, R package vegan). $R^2$: proportion of variance explained by each term. F: pseudo-F ratio. Treatment betadisper p: p-value of the multivariate dispersion homogeneity test between visitor treatments (betadisper/permutest, 9,999 permutations). For 16S, Bray-Curtis and weighted UniFrac $R^2$ values for CELESTO reflect near-complete dominance by a single Acinetobacter OTU in daytime-accessible samples.

| Genotype | Comparison_1 | n1 | Mean_1 | Comparison_2 | n2 | Mean_2 | *P*-value | *q-value* |
|---|---|---|---|---|---|---|---|---|
| CELESTO | No access | 7 | 1.46 ± 1.58 | Continuous access | 6 | 3.96 ± 0.10 | 0.011 | 0.032 |
| CELESTO | No access | 7 | 1.46 ± 1.58 | Daytime access | 7 | 4.00 ± 0.00 | 0.004 | 0.022 |
| CELESTO | No access | 7 | 1.46 ± 1.58 | Nighttime access | 6 | 2.54 ± 1.27 | 0.219 | 0.263 |
| CELESTO | Continuous access | 6 | 3.96 ± 0.10 | Daytime access | 7 | 4.00 ± 0.00 | 0.355 | 0.355 |
| CELESTO | Continuous access | 6 | 3.96 ± 0.10 | Nighttime access | 6 | 2.54 ± 1.27 | 0.06 | 0.09 |
| CELESTO | Daytime access | 7 | 4.00 ± 0.00 | Nighttime access | 6 | 2.54 ± 1.27 | 0.018 | 0.037 |
| IDILLIC | No access | 7 | 0.39 ± 0.32 | Continuous access | 3 | 1.00 ± 0.87 | 0.153 | 0.325 |
| IDILLIC | No access | 7 | 0.39 ± 0.32 | Daytime access | 3 | 1.58 ± 0.72 | 0.037 | 0.22 |
| IDILLIC | No access | 7 | 0.39 ± 0.32 | Nighttime access | 7 | 0.82 ± 0.67 | 0.217 | 0.325 |
| IDILLIC | Continuous access | 3 | 1.00 ± 0.87 | Daytime access | 3 | 1.58 ± 0.72 | 0.346 | 0.415 |
| IDILLIC | Continuous access | 3 | 1.00 ± 0.87 | Nighttime access | 7 | 0.82 ± 0.67 | 0.908 | 0.908 |
| IDILLIC | Daytime access | 3 | 1.58 ± 0.72 | Nighttime access | 7 | 0.82 ± 0.67 | 0.203 | 0.325 |

**Table S5. Pairwise Wilcoxon rank-sum test results for nectar microbial load (growth score) across pollinator access treatments, for CELESTO and IDILLIC cultivars.** Nectar was plated onto tryptic soy agar (TSA), R2A agar, potato dextrose agar (PDA) and yeast-malt (YM) agar, and incubated at 28 °C for 3–6 days (2024) or 4 days (2025). Growth score: semi-quantitative colony count per plate, scored 0 (no colonies), 1 (1–10), 2 (11–50), 3 (51–500), 4 (>500 colonies), see Figure S3. Tests were performed on per-sample means (mean score across the four culture media). n1, n2: number of samples per group; Mean_1, Mean_2: group mean ± SD of per-sample means. P-values were adjusted using the Benjamini–Hochberg procedure (six comparisons per cultivar × variable combination).

| Sample | Rep | Sucrose | Fructose | Glucose | Cultivar |
|---|---|---|---|---|---|
| 25ZM04_289 | R1 | -0.11 | 0.78 | 1 | CELESTO |
| 25ZM04_289 | R2 | -0.15 | 0.86 | 1 | CELESTO |
| 25ZM04_289 | R3 | 0.06 | 0.91 | 0.88 | CELESTO |
| 25ZM04_290 | R1 | -0.24 | NA | 1.94 | CELESTO |
| 25ZM04_290 | R2 | -0.17 | NA | 1.95 | CELESTO |
| 25ZM04_290 | R3 | -0.65 | 0.24 | 1.8 | CELESTO |
| 25ZM04_291 | R1 | -0.22 | 0.68 | 1.25 | CELESTO |
| 25ZM04_291 | R2 | 0.05 | 0.77 | 1.21 | CELESTO |
| 25ZM04_291 | R3 | -0.39 | 0.69 | 1.29 | CELESTO |
| 25ZM04_292 | R1 | -0.05 | 0.38 | 0.49 | CELESTO |
| 25ZM04_292 | R2 | -0.01 | 0.47 | 0.48 | CELESTO |
| 25ZM04_292 | R3 | 0.06 | 0.47 | 0.47 | CELESTO |
| 25ZM04_295 | R1 | 0.1 | 0.31 | 0.36 | IDILLIC |
| 25ZM04_295 | R2 | 0.38 | 0.28 | 0.25 | IDILLIC |
| 25ZM04_295 | R3 | 0.09 | 0.35 | 0.36 | IDILLIC |
| 25ZM04_296 | R1 | 0.22 | 0.18 | 0.21 | IDILLIC |
| 25ZM04_296 | R2 | 0.23 | 0.19 | 0.2 | IDILLIC |
| 25ZM04_296 | R3 | 0.2 | 0.18 | 0.21 | IDILLIC |
| 25ZM04_297 | R1 | 0.13 | 0.29 | 0.34 | IDILLIC |
| 25ZM04_297 | R2 | 0.05 | 0.21 | 0.45 | IDILLIC |
| 25ZM04_297 | R3 | 0.23 | 0.31 | 0.34 | IDILLIC |
| 25ZM04_298 | R1 | 0.23 | 0.23 | 0.24 | IDILLIC |
| 25ZM04_298 | R2 | 0.27 | 0.23 | 0.24 | IDILLIC |
| 25ZM04_298 | R3 | 0.23 | 0.22 | 0.24 | IDILLIC |

**Table S6. Enzymatically determined sucrose, fructose and glucose concentrations (g/L) in nectar, per technical replicate.** For each sample (one plant), nectar was pooled across 6 florets from the same head prior to three technical replicates (R1–R3). Data obtained with the Megazyme K-SUFRG assay. Negative sucrose values reflect assay readings near the detection limit. NA denotes a missing replicate measurement.

| OTU | Seq_Ref | Scientific Name | Query Cover | Identity |
|---|---|---|---|---|
| 77415788a2999e145f594d8f3738cd02e1e334f8 | ATTGAACGCTGGCGGCAGGCTTAACA-<br>CATGCAAGTCGAGCGGGGGAAGGTAGCTTGCTACTGGACCTAGCGGCGGACGG<br>GTGAGTAATACTTAGGAATCTGCCTATTAGTGGGGGACAACGTTCCGAAAGGAG<br>CGCTAATACCGCATACGCCCTACGGGGGAAAGCAGGGGATCTTCGGACCTTGCG<br>CTAATAGATGAGCCTAAGTCGGATTAGCTAGTTGGTGGGGTAAAGGCCTACCAA<br>GGCGACGATCTGTAGCGGGTTTGAGAGGATGATCCGCCACACTGGGACTGAGAC<br>ACGGCCCAGACTCCTACGGGAGGCAGCAGTGGGGAATATTGGACAATGGGCGC<br>AAGCCTGATCCAGCCATGCCGCGTGTGTGAAGAAGGCCTTATGGTTGTAAAGCAC<br>TTTAAGCGAGGAGGAGGCTTACCTAGCTAATATCTAGGCTAAGTGGACGTTACTC<br>GCAGAATAAGCACCGGCTAACTCTGTGCCAGCAGCCGCGGTAATACAGAGGGTG<br>CAAGCGTTAATCGGAATTACTGGGCGTAAAGCGCGCGTAGGTGGCCATTTAAGT<br>CAAATGTGAAATCCCCGAGCTTAACTTGGGAATTGCATTCGATACTGGATGGCTA<br>GAGTATAGGAGAGGAAGGTAGAATTCCAGGTGTAGCGGTGAAATGCGTAGAGA<br>TCTGGAGGAATACCGATGGCGAAGGCAGCCTTCTGGCCTAATACTGACACTGAG<br>GTGCGAAAGCATGGGGAGCAAACAGGATTAGATACCCTGGTAGTCCATGCCGTA<br>AACGATGTCTACTAGCCGTTGGGTCCTTTGAGGACTTAGTGGCGCAGCTAACGCG<br>ATAAGTAGACCGCCTGGGGAGTACGGTCGCAAGACTAAAACTCAAATGAATTGA<br>CGGGGGCCCGCACAAGCGGTGGAGCATGTGGTTTAATTCGATGCAACGCGAAGA<br>ACCTTACCTGGCCTTGACATACTAAGAACTTTCTAGAGATAGATTGGTGCCTTCGG<br>GAACTTAGATACAGGTGCTGCATGGCTGTCGTCAGCTCGTGTCGTGAGATGTTGG<br>GTTAAGTCCCGCAACGAGCGCAACCCTTTTCCTTACTTGCCAGCATTTCGGATGGG<br>AACTTTAAGGATACTGCCAGTGACAAACTGGAGGAAGGCGGGGACGACGTCAA<br>GTCATCATGGCCCTTACGGCCAGGGCTACACACGTGCTACAATGGTCGGTACAGA<br>GGGTTGCTACACAGCGATGTGATGCTAATCTCAAAAAGCCGATCGTAGTCCGGAT<br>TGGAGTCTGCAACTCGACTCCATGAAGTCGGAATCGCTAGTAATCGCGGATCAGA<br>ATGCCGCGGTGAATACGTTCCCGGGCCTTGTACACACCGCCCGTCACACCATGGG<br>AGTTTGTTGCACCAGAAGTAGATAGTCTAACCCTCGGGAGGACGTTTACCACGGT<br>GTGGCCAATGACTGGGGTG | *Acinetobacter nectaris* | 100% | 99.31% |
| 90c39638c0a1a1b6a2ca253c711811238359c694 | GATGAACGCTGGCGGCATGCTTAACACATGCAAGTCGGACGGGAAGTGG-<br>TGTTTCCAGTGGCGGACGGGTGAGTAACGCGTAAGAACCTGCCCTTGGGAGGGG<br>AACAACAGCTGGAAACGGCTGCTAATACCCCGTAGGCTGAGGAGCAAAAGGAG<br>GAATCCGCCCGAGGAGGGGCTCGCGTCTGATTAGCTAGTTGGTGAGGTAATAGC<br>TTACCAAGGCGATGATCAGTAGCTGGTCCGAGAGGATGATCAGCCACACTGGGA<br>CTGAGACACGGCCCAGACTCCTACGGGAGGCAGCAGTGGGGAATTTTCCGCAAT<br>GGGCGAAAGCCTGACGGAGCAATGCCGCGTGGAGGTAGAAGGCCCACGGGTCG<br>TGAACTTCTTTTCCCGGAGAAGAAGCAATGACGGTATCTGGGGAATAAGCATCG<br>GCTAACTCTGTGCCAGCAGCCGCGGTAATACAGAGGATGCAAGCGTTATCCGGA<br>ATGATTGGGCGTAAAGCGTCTGTAGGTGGCTTTTTAAGTCCGCCGTCAAATCCCA<br>GGGCTCAACTCTGGACAGGCGGTGGAAACTACCAAGCTGGAGTACGGTAGGGG<br>CAGAGGGAATTTCCGGTGGAGCGGTGAAATGCGCAGAGATCGGAAAGAACACC<br>AACGGCGAAAGCACTCTGCTGGGCCGACACTGACACTGAGAGACGAAAGCTAGG<br>GGAGCGAATGGGATTAGATACCCCAGTAGTCCTAGCCGTAAACGATGGATACTA<br>GGCGCTGTGCGTATCGACCCGTGCAGTGCTGTAGCTAACGCGTTAAGTATCCCGC<br>CTGGGGAGTACGTTCGCAAGAATGAAACTCAAAGGAATTGACGGGGGCCCGCAC<br>AAGCGGTGGAGCATGTGGTTTAATTCGATGCAAAGCGAAGAACCTTACCAGGGC<br>TTGACATGCCGCGAATCCTCTTGAAAGAGAGGGGTGCCTTCGGGAACGCGGACA<br>CAGGTGGTGCATGGCTGTCGTCAGCTCGTGCCGTAAGGTGTTGGGTTAAGTCCC<br>GCAACGAGCGCAACCCTCGTGTTTAGTTGCCATCATTGAGTTTGGAACCCTGAAC<br>AGACTGCCGGTGATAAGCCGGAGGAAGGTGAGGATGACGTCAAGTCATCATGCC<br>CCTTATGCCCTGGGCGACACACGTGCTACAATGGCCGGGACAAAGGGTCGCGAT<br>CCCGCGAGGGTGAGCTAACTCCAAAAACCCGTCCTCAGTTCGGATTGCAGGCTGC<br>AACTCGCCTGCATGAAGCCGGAATCGCTAGTAATCGCCGGTCAGCCATACGGCG<br>GTGAATCCGTTCCCGGGCCTTGTACACACCGCCCGTCACACTATGGGAGCTGGCC<br>ATGCCCGAAGTCGTTACCTTAACCGCAAGGAGGGGGATGCCGAAGGCAGGGCTA<br>GTGACTGGAGTG | *Helianthus annuus* | 100% | 100% |

**Table S7. BLAST alignment results for the two dominant 16S OTUs.** The *Acinetobacter* OTU was queried against the NCBI 16S ribosomal RNA sequences database (Bacteria and Archaea), the chloroplast OTU was queried against the NCBI nucleotide database (nr/nt), as plastidial sequences are not represented in the bacterial 16S database. Query cover and percent identity refer to the top BLASTn hit. Accession numbers for top hits: *Acinetobacter nectaris* NR_118408.1; *Helianthus annuus* MK341449.1.

| Compound | Annotation | Cultivar | No access | Continuous access | Daytime access | Nighttime access | Model P | Model q |
|---|---|---|---|---|---|---|---|---|
| VOC_6262 | alpha-Guaiene | CELESTO | a | b | a | a | 3.55E-04 | 0.0113 |
| VOC_6763 | alpha-Maaliene | CELESTO | a | b | a | a | 2.89E-04 | 0.0113 |
| VOC_6915 | Unidentified compound | CELESTO | a | b | a | a | 4.06E-04 | 0.0113 |
| VOC_8514 | Maalilol | CELESTO | a | b | a | a | 3.32E-05 | 0.00368 |
| VOC_2218 | Unidentified compound | IDILLIC | a | b | a | a | 0.00232 | 0.0288 |
| VOC_3669 | Not checked | IDILLIC | ab | a | b | b | 0.00233 | 0.0288 |
| VOC_3760 | Not checked | IDILLIC | a | a | b | b | 1.12E-04 | 0.00619 |
| VOC_4573 | Verbenone | IDILLIC | a | b | a | a | 0.00232 | 0.0288 |
| VOC_6553 | alpha-Guaiene like | IDILLIC | a | b | ab | a | 0.00334 | 0.0371 |
| VOC_6700 | Calarene (beta-Gurjunene) | IDILLIC | a | b | ab | a | 0.00222 | 0.0288 |
| VOC_6763 | alpha-Maaliene | IDILLIC | a | b | ab | a | 7.66E-04 | 0.0213 |
| VOC_6915 | Unidentified compound | IDILLIC | a | b | a | a | 1.11E-04 | 0.00619 |
| VOC_8514 | Maalilol | IDILLIC | a | b | a | a | 3.20E-04 | 0.0118 |
| VOC_8627 | ((Iso-)Spathulenol) | IDILLIC | a | b | a | a | 0.00155 | 0.0288 |

**Table S8. Volatile compounds significantly affected by pollinator access treatment (Benjamini–Hochberg adjusted model q < 0.05), by cultivar.** Log10(x + 1) transformed abundance was modelled against the four-level access treatment factor separately for each cultivar (ANOVA). Correction was applied within each cultivar across the 111 compounds tested. For compounds passing this threshold, each cell gives the Šidák-adjusted compact letter grouping for that access modality. Modalities sharing a letter within a compound do not differ significantly from one another. Model P and q are the raw and Benjamini–Hochberg-adjusted P-values for the overall treatment effect. Annotation status: named compounds were confirmed by manual inspection of raw spectral data (library match plus expert curation). "Unidentified compound" indicates a genuine spectral feature that could not be matched to a library reference. "Not checked" indicates manual curation of the annotation against raw data has not yet been performed.
